# A Mouse Model of Brittle Cornea Syndrome caused by mutation in *Zfp469*

**DOI:** 10.1101/2021.07.08.451591

**Authors:** Chloe M. Stanton, Amy S. Findlay, Camilla Drake, Mohammad Z. Mustafa, Philippe Gautier, Lisa McKie, Ian J. Jackson, Veronique Vitart

**Affiliations:** MRC Human Genetics Unit, Institute of Genetics & Cancer, The University of Edinburgh, Western General Hospital, Crewe Road, Edinburgh, EH4 2XU

**Keywords:** Brittle cornea, keratocyte, Zfp469, ZNF469, collagen

## Abstract

Brittle Cornea Syndrome (BCS) is a rare recessive condition characterised by extreme thinning of the cornea and sclera. BCS results from loss-of-function mutations in the poorly understood genes *ZNF469* or *PRDM5*. In order to determine the function of ZNF469 and to elucidate pathogenic mechanisms, we used genome editing to recapitulate a human *ZNF469* BCS mutation in the orthologous mouse gene, *Zfp469*. Ophthalmic phenotyping showed that homozygous Zfp469 mutation causes significant central and peripheral corneal thinning arising from reduced stromal thickness. Expression of key components of the corneal stroma in primary keratocytes from *Zfp469*^BCS/BCS^ mice is affected, including decreased *Col1a1* and *Col1a2* expression. This alters the type I:type V collagen ratio and results in collagen fibrils with smaller diameter and increased fibril density in homozygous mutant corneas, correlating with decreased biomechanical strength in the cornea. Cell-derived matrices generated by primary keratocytes show reduced deposition of type I collagen offering an *in vitro* model for stromal dysfunction. Work remains to determine whether modulating ZNF469 activity will have therapeutic benefit in BCS or in conditions such as keratoconus where the cornea thins progressively.

**Summary statement:** A mouse model of Brittle Cornea Syndrome was created to elucidate molecular mechanisms underlying pathology of this rare connective tissue disorder in which extremely thin corneas rupture, causing irreversible blindness.

## Introduction

Brittle Cornea Syndrome (BCS; MIM 229200, MIM 614170) is a rare autosomal recessive disorder that is characterised by extreme thinning of the cornea and sclera. Visual impairment may initially be a result of myopia and progressive keratoconus or keratoglobus but, as the name suggests, the thin and fragile corneas of affected individuals are prone to rupture leading to irreversible blindness (Al-Hussain et al., 2004). Also classified as a subtype of Ehlers-Danlos syndrome (EDS type VIB), this devastating condition often leads to general connective tissue dysfunction with skin hyperelasticity, joint hyperflexibility and, in approximately one third of cases, hearing impairment (Malfait et al., 2017).

BCS results from biallelic loss-of-function (LOF) mutations in *ZNF469* or *PRDM5* (Abu et al., 2008; Burkitt Wright et al., 2011). Mutations in these genes appear to cause an indistinguishable disorder, suggesting that they contribute to the same biological pathways. PRDM5 (PR/SET Domain 5), a widely expressed transcription factor that modulates development and maintenance (Duan et al., 2007; Meani et al., 2009; Porter et al., 2015) is known to play an important role in extracellular matrix (ECM) production by several tissues including skin fibroblasts (Porter et al., 2015) and in bone (Galli et al., 2012). The role played by *ZNF469* in the healthy cornea and in BCS is less clear as the function of the very large protein encoded by this gene is poorly characterised. Since mutations in *ZNF469* were first reported to cause BCS (Abu et al., 2008), 30 pathogenic compound heterozygous or homozygous mutations have been identified in the single coding exon of *ZNF469* spanning 13 kilobases on chromosome 16q24.2 (Abu et al., 2008; Al-Owain et al., 2012; Christensen et al., 2010; Dhooge et al., 2021; Khan et al., 2012; Khan et al., 2010; Menzel-Severing et al., 2019; Micheal et al., 2019; Ramappa et al., 2014; Rohrbach et al., 2013; Rolvien et al., 2020; Skalicka et al., 2020). The majority of these mutations result in premature stop codons that are hypothesised to lead to production of truncated and non-functional protein (Rohrbach et al., 2013) and are thus considered as loss of function (LOF) mutations.

Since *ZNF469* encodes a C2H2 zinc-finger (ZF) protein, it is assumed to function as a transcriptional regulator for the synthesis or assembly of collagen fibrils (Abu et al., 2008). Studies in dermal fibroblasts showed that mutations in ZNF469 or PRDM5 both affect the expression of fibrillar collagen genes and the deposition of collagen into the extracellular matrix (ECM) (Burkitt Wright et al., 2011). In humans, 90% of corneal thickness (normal range 450–700 μm (Vitart et al., 2010)) is contributed by the stroma (Alberto and Garello, 2013), a collagen-rich ECM deposited by resident keratocytes. The precise organisation of collagen fibrils in the stroma is crucial to cornea function, providing approximately two thirds of the human eye’s refractive power whilst combining mechanical strength and almost perfect transmission of visible light. Stromal disorganisation and thinning are seen in many corneal disorders, including keratoconus (affecting 1/2000 individuals in the UK (Pearson et al., 2000)), but the reduction in CCT observed in BCS is extreme (<400 μm).

The underlying biological processes that lead to corneal thinning in BCS are not well understood, but may reflect pathological extremes of the processes determining corneal thickness in healthy eyes. This is supported by heterozygous carriers of *ZNF469* and *PRDM5* mutations, who have mildly reduced central corneal thickness (CCT) relative to homozygotes (Burkitt Wright et al., 2011), and by the association of variants in putative regulatory elements of *ZNF469* with reduced CCT in the general population (Iglesias et al., 2018; Lu et al., 2010; Lu et al., 2013). Dysregulation of collagen synthesis or assembly by keratocytes is known to have a strong effect on corneal thickness (Dimasi et al., 2010; Evereklioglu et al., 2002; Segev et al., 2006; Sun et al., 2011). However, the role that ZNF469 plays in keratocyte cells remains undetermined. In order to address this, we generated the first mouse model of BCS caused by mutation in the orthologous mouse gene *Zfp469* to elucidate the mechanisms by which stromal thickness is controlled in health and disease.

## Results

### Zfp469 is the mouse ortholog of human ZNF469 and can be modified to recapitulate human BCS mutations

We performed a literature review to compile a comprehensive list of mutations in *ZNF469* that have been identified in BCS families to date (March 2021), and mapped each mutation to human *ZNF469* transcript NM_001367624.2 (Table 1). Of the 30 genetic variants that have been reported as homozygous or compound heterozygous mutations in BCS cases, 27 (90%) result in either nonsense or frameshift mutations in ZNF469 predicted to result in production of a truncated protein. These mutations occur throughout the *ZNF469* coding sequence (NP_001354553.1) (Fig. 1A), and, with the exception of one large deletion encompassing the gene (Ramappa et al., 2014), share common consequences of truncating the protein or, in the case of two missense mutations, to modify key residues involved in the coordination of zinc in the sixth C2H2 ZF domain. Fewer LOF mutations than expected have been seen in human populations (probability of being loss-of-function intolerant, pLI, = 0.72 (observed/expected = 0.2 (0.12 – 0.37, 90% Confidence Interval) in gnomAD (Lek et al., 2016). Only four of the BCS mutations reported in affected individuals, p.Gln2149Serfs*51, p.Gln2149Alafs*42, p.Arg3442Glyfs*59 and p.Pro3584Glnfs*136, have been identified in a small number of heterozygous individuals in gnomAD v3.1 (Table 1).

**Fig. 1.**
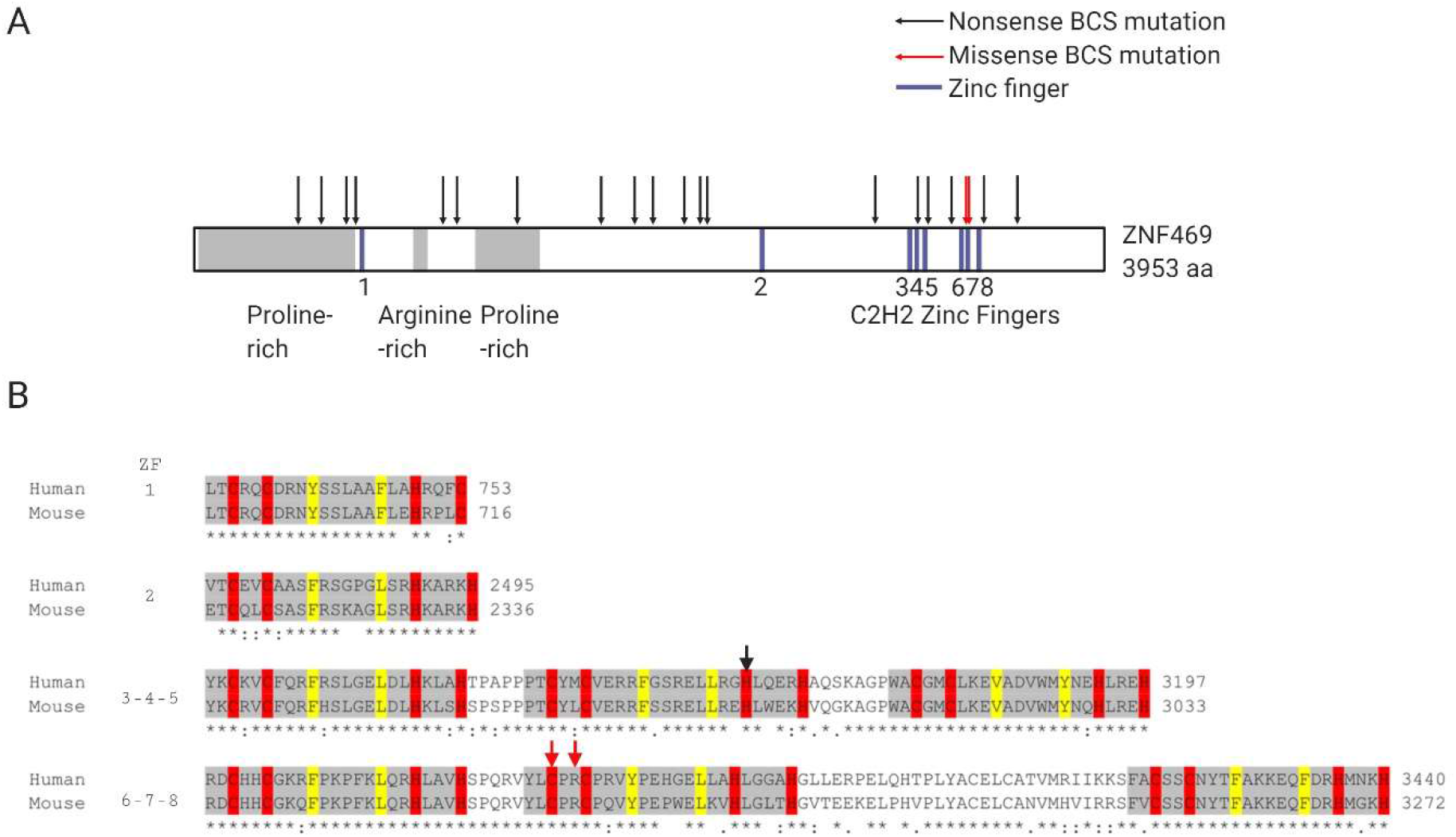
Pathogenic mutations in ZNF469 are distributed throughout the protein and result in premature termination codons or disruption of conserved residues in C2H2 zinc finger domains. (A) Schematic representation of the protein structure of ZNF469, showing the position of previously reported pathogenic variants in relation to 8 C2H2 zinc finger domains and regions of compositional bias. Nonsense or frameshift variants resulting in a premature stop are indicated by a black arrow; missense mutations are indicated by a red arrow. (B) Sequences of the 8 C2H2 zinc finger domains in the mouse (Zfp469, NP_001354553.1) and human (ZNF469, NP_001354553.1) proteins were aligned using Clustal Omega. Fully conserved residues are indicated by *, : indicates conservation between groups of amino acids with strongly similar properties, and . indicates conservation between groups of amino acids with weakly similar properties. Residues shown in red are the paired cysteines (C) and histidines (H) bind the zinc ion. Residues in yellow are structurally important hydrophobic amino acids.

**Table 1.** Pathogenic mutations in *ZNF469* previously reported in Brittle Cornea Syndrome.

| Human Chromosomal location (GRCh38/hg38) | Transcript NM_001367624.2 | NP_001354553.1 Amino acid change | Consequence | Variant ID | Allele Frequency in gnomAD v3.1 | SIFT | PolyPhen | CADD | Amino acid Conservation in mouse | Reference |
| --- | --- | --- | --- | --- | --- | --- | --- | --- | --- | --- |
| 16:88438217-88438218 | c.10748delC | p.Pro3584Glnfs*136 | frameshift | rs1253064545, COSV71258333 | 0.00000658 | - | - | 24.8 | not conserved | Rolvien et al., 2020 |
| 16:88437793-88437794 | c.10324delA | p.Arg3442Glyfs*59 | frameshift | rs756543273 | 0.00003942 | - | - | 22.6 | not conserved | Rolvien et al., 2020 |
| 16:88437576-88437576 | c.10106G>C | p.Arg3369Pro | missense | CM134610 | - | Deleterious (0.01) | Probably damaging (0.935) | 25.1 | yes | Rohrbach et al., 2013 |
| 16:88437570-88437570 | c.10100G>A | p.Cys3367Tyr | missense | rs387907062, CM100171 | - | Deleterious (0) | Probably damaging (1) | 33 | yes | Christensen et al., 2010 |
| 16:88437385-88437385 | c.9915dupC | p.Arg3306Glnfs*197 | frameshift | - | - | - | - | - | not conserved | Micheal et al., 2019 |
| 16:88437346-88437346 | c.9876dupT | p.Ala3293Cysfs*6 | frameshift | - | - | - | - | - | not conserved | Dhooge et al., 2021 |
| 16:88437080-88437081 | c.9611delG | p.Gln3206Argfs*23 | frameshift | - | - | - | - | 23.3 | yes | Abu et al., 2008 |
| 16:88436952-88436953 | c.9483delG | p.His3162Thrfs*20 | frameshift | CD134609 | - | - | - | - | yes | Rohrbach et al., 2013 |
| 16:88436384-88436384 | c.8901_8914dup | p.Glu2972Glyfs*50 | frameshift | - | - | - | - | - | not conserved | Al-Owain et al., 2012 |
| 16:88434197-88434198 | c.6728delA | p.Asp2243Alafs*8 | frameshift | rs1163546262, CD1414349 | - | - | - | 18.76 | not conserved | Ramappa et al., 2014 |
|  | deletion of 16q24.1 | Gene deletion |  | - | - | - | - | - | - | Ramappa et al., 2014 |
| 16:88434116-88434117 | c.6647delA | p.Gln2216Argfs*19 | frameshift | CD134604 | - | - | - | 16.86 | not conserved | Rohrbach et al., 2013 |
| 16:88433913-88433914 | c.6444delG | p.Gln2149Serfs*51 | frameshift | CD134602, COSV71259221 | 0.00000657 | - | - | - | not conserved | Rohrbach et al., 2013 |
| 16:88433913-88433914 | c.6444dupG | p.Gln2149Alafs*42 | frameshift | COSV101458941 | 0.00000657 |  |  | - | not conserved | Menzel-Severing et al., 2019 |
| 16:88433496-88433497 | c.6027delA | p.Gly2011Alafs*16 | frameshift | CD082220 | - | - | - | 19.02 | not conserved | Abu et al., 2008 |
| 16:88433257-88433258 | c.5788dupC | p.Gln1930Profs*133 | frameshift | - | - | - | - | - | not conserved | Rohrbach et al., 2013 |
| 16:88433258-88433258 | c.5788delC | p.Gln1930Argfs*6 | frameshift | CD134600 | - | - | - | - | not conserved | Rohrbach et al., 2013 |
| 16:88432823-88432823 | c.5353C>T | p.Gln1785* | stop_gained | CM1111667 | - | - | - | 34 | not conserved | Burkitt Wright et al., 2011 |
| 16:88431728-88431728 | c.4258G>T | p.Glu1420* | stop_gained | rs387907063, CM106782 | - | - | - | 34 | not conserved | Khan et al., 2010 |
| 16:88430945-88430946 | c.3476delG | p.Gly1159Alafs*105 | frameshift | CD134598 | - | - | - | - | not conserved | Rohrbach et al., 2013 |
| 16:88430777-88430777 | c.3307C>T | p.Gln1103* | stop_gained | - | - | - | - | 33 | not conserved | Dhooge et al., 2021 |
| 16:88430774-88430774 | c.3304G>T | p.Glu1102* | stop_gained | - | - | - | - | 35 | not conserved | Rohrbach et al., 2013 |
| 16:88429619-88429620 | c.2150delT | p.Phe717Serfs*15 | frameshift | CD129552 | - | - | - | - | yes | Khan et al., 2012 |
| 16:88429499-88429499 | c.2029G>T | p.Gly677* | stop_gained | CM134439 | - | - | - | 38 | yes | Rohrbach et al., 2013 |
| 16:88429433-88429433 | c.1963dupC | p.His655Profs*83 | frameshift | COSV71259490 | - | - | - | - | not conserved | Menzel-Severing et al., 2019 |
| 16:88429175-88429175 | c.1705C>T | p.Gln569* | stop_gained | COSV71258665 | - | - | - | 37 | yes | Skalicka et al., 2019 |
| 16:88429055-88429056 | c.1586delG | p.Gly529Aspfs*9 | frameshift | - | - | - | - | - | not conserved | Dhooge et al., 2021 |
| 16:88428913-88428914 | c.1444delC | p.Leu482Cysfs*20 | frameshift | COSV71258360 | - | - | - | - | yes | Dhooge et al., 2021 |
| 16:88428871-88428881 | c.1402_1411del | p.Pro468Alafs*31 | frameshift | - | - | - | - | - | not conserved | Skalicka et al., 2019 |
| 16:88428550-88428551 | c.1081delG | p.Ala361Leufs*16 | frameshift | - | - | - | - | - | yes | Dhooge et al., 2021 |

The orthologous gene in mouse, *Zfp469* or *Gm22,* is located on chromosome 8 (NM_001362883). Clustal Omega (Sievers et al., 2011) was used to align orthologous protein sequences from mouse (Zfp469, NP_001354553.1) and human (ZNF469, NP_001354553.1), with 47% amino acid identity across the full-length of each protein (human 3953 amino acids, mouse 3765 amino acids). The 8 predicted C2H2 ZF domains (positions obtained from the SMART database (Letunic et al., 2020)) show a much higher degree of conservation (Fig.1B), consistent with ZF domains having an important functional role. Of the human BCS mutations, only some affected conserved amino acids (Table 1). One of these (human p.Gly677*, mouse p.Gly634) was selected as the target for genome editing using CRISPR-Cas9n to create a mouse model of BCS.

### Gene editing *Zfp469* to generate *Zfp469*^BCS^ mice

In order to elucidate the role of Zfp469 in mice, genome editing was performed to recapitulate a human BCS mutation by creating a premature stop codon early in *Zfp469* the C2H2 ZF domains. Three pairs of sgRNA (Table S1) designed to target sequence encoding Zpf469 p.Gly634 were cloned into pX458 (pSpCas9(BB)-2A-GFP) (Ran et al., 2013) and tested for cleavage efficiency in mouse embryonic fibroblasts (MEFs). The pair of sgRNAs with the highest cleavage efficiency (data not shown) was subsequently *in vitro* transcribed, and purified RNA was injected, along with SpCas9n mRNA and the ssODN repair template, into C57Bl/6J mouse zygotes. Successful genome editing and repair inserted an in-frame V5 tag and generated a premature stop codon to truncate Zfp469 (Fig.2A). Homozygous mutant mice were significantly smaller than their sex-matched litter-mate controls at 3 months of age (Fig.2B and 2C), weighing 15-20% less than wildtype mice (males, p=0.003, females p=0.0429, one way ANOVA with Dunnett’s multiple comparison test). Body length was decreased by approximately 5% in *Zfp469*^BCS/BCS^ mice relative to wildtype age- and sex-matched littermates, but this difference was not statistically significant at 3 or 6 months of age (data not shown). There was no difference in eye size, measured using manual calipers, between genotypes in females at 3 months of age (average eye length from the front of the cornea to the posterior: ^+/+^ 3.34 ± 0.13 mm (s.d., n=3), ^BCS/BCS^ 3.33± 0.13 mm (s.d., n=2). Other than the reduction in body weight, heterozygous and homozygous *Zfp469*^BCS^ mice were viable and fertile, with each genotype arising in the expected frequencies from heterozygote × heterozygote crosses (Table S2).

**Fig. 2.**
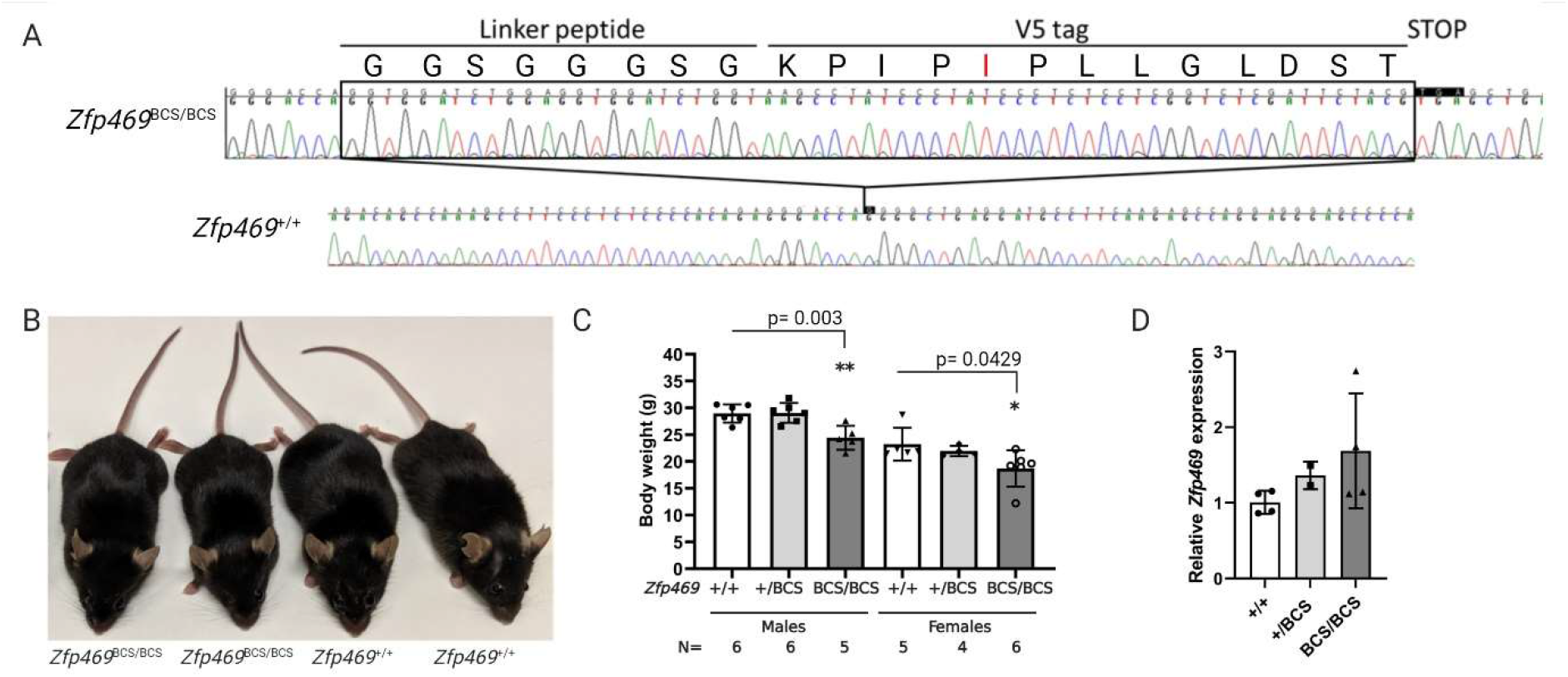
Genome editing *Zfp469* to create a mouse model of Brittle Cornea Syndrome (BCS). (A) CRISPR-Cas9 genome editing was used to insert an in-frame V5 tag and premature stop codon into Zfp469 at p.Gly634, the conserved residue equivalent to the human mutation p.Gly677*. Sequencing chromatograms from a homozygous *Zfp469*^BCS/BCS^ mouse compared to a wildtype litter mate indicating the position and sequence of the insertion. The V5 tag contained a spontaneously arising A>T mutation, resulting in the amino acid change Asn (N) >Ile (I) at position 5 in the V5 tag. (B) A representative image of 2 female *Zfp469*^BCS/BCS^ and 2 female *Zfp469*^+/+^ littermate mice at 3 months of age. *Zfp469*^BCS/BCS^ mice show no gross phenotypic abnormalities but are smaller than *Zfp469*^+/+^ sex-matched littermates. (C) Body weight of *Zfp469*^BCS/BC^ is significantly reduced relative to *Zfp469*^+/+^ age- and sex-matched animals, as determined by one way ANOVA with Dunnett’s multiple comparison test. Data are presented as mean ± s.d.. (D) qPCR analysis revealed no significant difference in the relative expression of *Zfp469* in keratocytes isolated from the corneas of 4 wildtype, 2 heterozygous and 4 homozygous mice aged 3 months. Data are presented as mean ± s.d. with the average of assays performed in duplicate from each sample shown by data points.

Expression of *Zfp469* in corneal keratocytes freshly isolated from corneas pooled by genotype was assessed by RT-qPCR using primers specific for *Zfp469* or for the V5-tag insertion. Relative *Zfp469* expression was increased to 1.69 ± 0.64 (s.d.) in homozygotes (1.00 ± 0.15 (s.d.) in wildtypes) but was not significantly different between genotypes (Fig. 2D). All of the *Zfp469* transcript expressed in homozygous mutant and 55% ± 3.7% (s.d.) of transcript in heterozygous keratocytes contained the V5 sequence showing that the mutant transcript is expressed at a similar level to the wildtype transcript. This is consistent with a premature stop codon in the single coding exon of Zfp469 resulting in the production of a truncated protein rather than destruction of the transcript by nonsense-mediated decay (NMD) (Fig.2D).

### *Zfp469*^BCS^ mice recapitulate ophthalmic characteristics of BCS

Clinical characteristics of BCS include thinning of the cornea and sclera, myopia and refractive errors, keratoconus or keratoglobus, corneal rupture and vision loss. We performed ophthalmic phenotyping including slit lamp examination, anterior-segment optical coherence tomography (AS-OCT), histology and immunostaining in order to determine whether the premature stop codon introduced into *Zfp469* leads to features of BCS in the mouse. At 3 months of age, by which stage the cornea is fully developed (Hanlon et al., 2011), slit lamp examination revealed no gross corneal opacity or abnormality in homozygous mice (Fig.3A). However, AS-OCT (Fig.3B) showed that homozygous mice have extremely thin corneas relative to age- and sex-matched wildtype animals, with a mean reduction of 39.1 ± 4.4 μm (s.e.m.) in central corneal thickness (p = 0.0025 in males, p<0.0001 in females, p<0.0001 for sexes combined, one way ANOVA with Tukey’s multiple comparison test) (Fig. 3C). This corresponds to a 30% reduction in CCT. Thickness of the peripheral corneal was also significantly reduced by approximately 25% (in female homozygous mice, average 144.6 ± 15.3 μm s.d., n=4; in wildtype females, 187.9 ± 16.2 μm s.d., n=3, p = 0.0335) consistent with generalised thinning of the cornea as is seen in BCS.

**Fig. 3.**
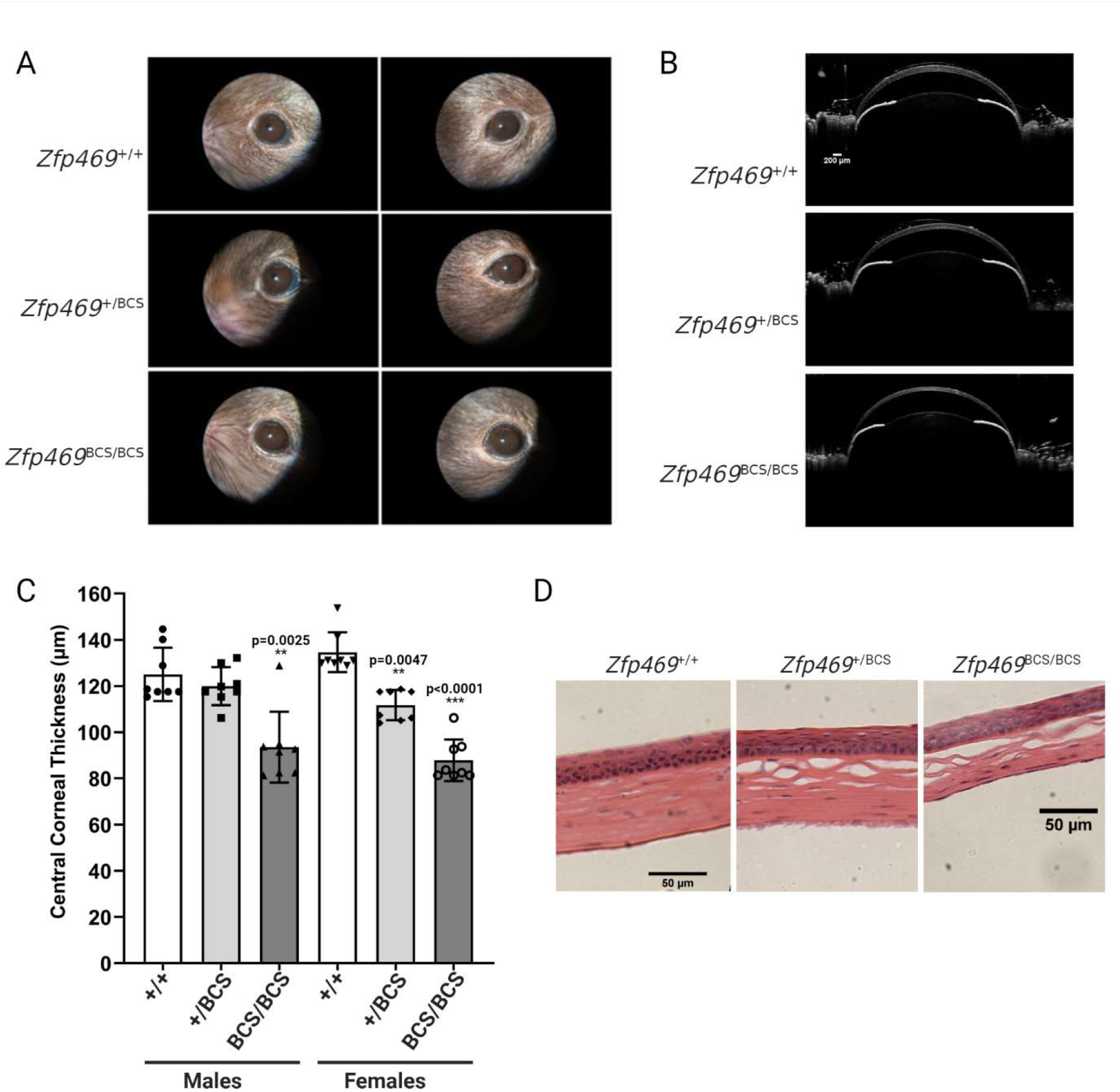
Corneal thinning in *Zfp469*^BCS/BCS^ mice as a result of decreased stromal thickness. (A) Corneal opacity is not observed in 3 month old *Zfp469*^BCS/BCS^ mice examined by slit lamp. (B) Anterior-segment optical coherence tomography (AS-OCT) of 3 month old *Zfp469*^BCS/BCS^ mice demonstrates extreme thinning of the central and peripheral cornea compared to *Zfp469*^+/+^ and *Zfp469*^+/BCS^ age- and sex-matched mice. Scale bar = 200 μm. (C) Central Corneal thickness (CCT) is decreased in *Zfp469*^BCS/BCS^ mice (n=4 for each genotype and sex; p=0.0025 in males, p<0.0001 in females, p<0.0001 for sexes combined, one way ANOVA with Tukey’s multiple comparison test). Data are presented as mean ± s.d., with individual measurements from each eye shown by data points. (D) Heamatoxylin and eosin staining of corneal sections from 3 month old mice reveals visibly thinner corneal stroma in *Zfp469*^BCS/BCS^ mice. Scale bar is 50 μm.

Hematoxylin and eosin (H&E) staining to examine corneal morphology confirmed that the multi-layered epithelium appeared normal, as does the intact endothelium (Fig.3D). However, the corneal stroma, which normally comprises 90% thickness of a healthy cornea in humans and approximately 60% of thickness in mice (Hanlon et al., 2011), is markedly thinner in homozygotes. Extensive shearing was observed between lamellae relative to that seen in wildtype cornea processed for staining in the same way.

### Corneal thinning in *Zfp469*^BCS^ mice is apparent during corneal development and is not progressive

In some clinical reports, the corneal thinning observed in BCS patients is described as progressive and corresponds with an increasing risk of spontaneous rupture of the cornea or sclera. We sought to determine whether the corneal thinning observed in the *Zfp469*^BCS/BCS^ mice at 3 months was established earlier, during development of the cornea, and whether thinning of the stroma was progressive. At 1 month of age, AS-OCT (Fig.4A) revealed that the central corneas of homozygous animals were, on average, already 46.8 ± 5.6 μm (s.e.m.) thinner than wildtype sex- and age-matched controls (Fig.4B, p = 0.0016 males, p = 0.0043 females, p<0.0001 for sexes combined, one way ANOVA with Tukey’s multiple comparison test) - a 35% reduction in CCT. Fig.4C shows a unilateral corneal opacity observed in one eye from a *Zfp469*^BCS/BCS^ female at 1 month of age (n=12 studied). OCT revealed swelling of the corneal stroma consistent with an edema. It was not possible to determine whether this was a result of injury leading to rupture of Descemet’s membrane but there was no evidence of injury or infection. No such accumulation of fluid within the cornea was observed in 26 heterozygous or wildtype eyes at 1 month of age, or in any older mice.

**Fig. 4.**
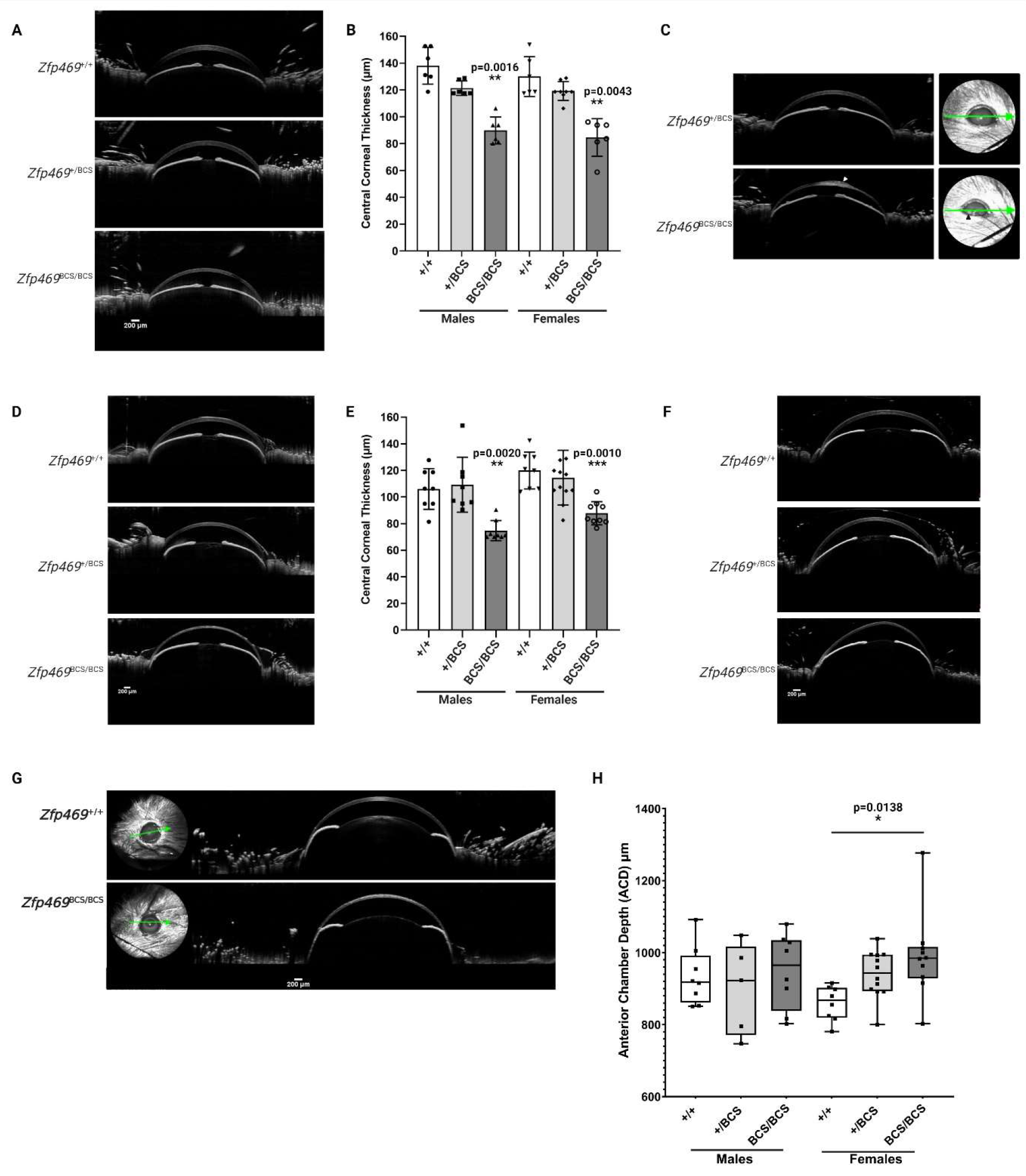
Corneal thinning in *Zfp469*^BCS^ mice is apparent during corneal development and is not progressive. (A) Extreme thinning of the cornea in 1 month old *Zfp469*^BCS/BCS^ mice compared to *Zfp469*^+/+^ and *Zfp469*^+/BCS^ age- and sex-matched mice is observed by AS-OCT Scale bar is 200 μm. (B) At 1 month of age, CCT is significantly decreased in *Zfp469*^BCS/BCS^ mice (p=0.0016 in males, p=0.0043 in females, p<0.0001 for sexes combined, one way ANOVA with Tukey’s multiple comparison test). N=3 for each genotype in males; n=3 for *Zfp469*^+/+^ and *Zfp469*^BCS/BCS^, n=4 for *Zfp469*^+/BCS^ in females. Data are presented as mean ± s.d., with individual measurements from each eye shown by data points. (C) Unilateral corneal opacity observed in one eye from a *Zfp469*^BCS/BCS^ female at 1 month of age resembles corneal edema (white arrowhead). (D) CCT at 6 months of age remains similar to CCT measured at 1 and 3 months of age, with significant central and peripheral thinning apparent only in *Zfp469*^BCS/BCS^ mice. Scale bar is 200 μm. (E) CCT is significantly decreased in *Zfp469*^BCS/BCS^ mice at 6 months of age (p=0.0020 in males, p=0.0010 in females, p<0.0001 for sexes combined, one way ANOVA with Tukey’s multiple comparison test). N=4 for each genotype in males; n=4 for *Zfp469*^+/+^, n=6 for *Zfp469*^+/BCS^, n=5 for *Zfp469*^BCS/BCS^ in females. Data are presented as mean ± s.d., with individual measurements from each eye shown by data points. (F) Corneal distortion is seen by AS-OCT after application of eye drops in *Zfp469*^BCS/BCS^ eyes, but not in heterozygous or wildtype sex-matched littermates, suggesting a loss of biomechanical strength in the mutant corneas. Scale bar is 200 μm. (G) Globular protrusion of the eye resembling keratoglobus is observed after dilation of the pupil in *Zfp469*^BCS/BCS^ female mice, but not in male mice or heterozygous or wildtype sex-matched controls. Scale bar is 200 μm. (H) Anterior chamber depth (ACD) in the eyes of *Zfp469*^BCS/BCS^ female mice, measured from AS-OCT images, is significantly increased compared to heterozygous or wildtype sex-matched controls after dilation of the pupil. p=0.0047, Welch’s ANOVA test). n=4 for *Zfp469*^+/+^, n=3 for *Zfp469*^+/BCS^, n=4 for *Zfp469*^BCS/BCS^ in males; n=4 for *Zfp469*^+/+^, n=6 for *Zfp469*^+/BCS^, n=5 for *Zfp469*^BCS/BCS^ in females. Box plot shows all data, with individual measurements from each eye shown by data points. Whiskers show minimum to maximum values.

Subsequent investigation of corneal thickness at 6 months of age also showed a 30% reduction in CCT, with mean CCT 31.4 ± 5.2 μm (s.e.m.) thinner in *Zfp469*^BCS/BCS^ (Fig.4D, E, p=0.0023 males, p=0.001 females, p<0.0001 combined). Peripheral corneal thickness was also significantly reduced (in female homozygous mice, average 125.99 ± 14.1 μm, s.d., n=4; in wildtype females, 166.9 ± 21.8 μm s.d., n=3, p = 0.0286). The degree of thinning observed at 1, 3 and 6 months consistently reduced CCT by approximately 30% relative to wildtype age- and sex-matched mice. The corneas of individual mice with homozygous BCS mutation in Zfp469 - followed from 3 months of age - do not progressively thin up to age 6 months.

The thin corneas of homozygous mutant mice aged up to 9 months were not prone to perforation; no animals were affected by corneal damage under normal housing conditions. However, the *Zfp469*^BCS/BCS^ corneas were more easily distorted by application of a viscous liquid drop intended to prevent the eye from drying out during OCT than wildtype controls at both 3 and 6 months of age (Fig.4F). We also measured anterior chamber depth (ACD) in AS-OCT images obtained of female mice at 6 months of age. Without dilation of the pupils, ACD in *Zfp469*^+/+^ was 977.1 μm ± 32.9 (mean ± s.d., n=3), in *Zfp469*^+/BCS^ 1002.9 μm ± 55.1 (mean ± s.d., n=5), and in *Zfp469*^BCS/BCS^ 1016.5 μm ± 32.1 (mean ± s.d., n=4). This difference was not significant but suggested there may be a trend towards increased ACD in eyes of homozygous mutant mice. Subsequently, when the pupils of the same animals were dilated by administration of Tropicamide and Phenylephrine Hydrochloride prior to performing AS-OCT, female *Zfp469*^BCS/BCS^ showed a bulging eye phenotype consistent with keratoglobus (Fig.4G), a condition in which generalised corneal thinning results in globular protrusion of the eye. This was not observed in male mice, or in wildtype females. ACD was determined from OCT images after pupil dilation, confirming a significant increase in ACD in *Zfp469*^BCS/BCS^ females relative to wildtype controls at 6 months of age (Fig.4H, Welch’s ANOVA test, p=0.0138). Interestingly, the ACD of homozygous mutant eyes decreased by only 2.6% after dilation of the pupil. This contrasts to wildtype ACD which was reduced by 12% on average relative to ACD determined without dilation of the pupil. Together with the specific thinning of the stroma revealed by H&E staining, these data are indicative of a loss of biomechanical strength of the corneal stroma resulting in impaired resistance to both internal and external forces.

### Collagen type I, but not type V, is less abundant in the *Zfp469*^BCS/BCS^ cornea and fibril architecture is altered

The corneal stroma is a collagenous extracellular matrix generated by resident keratocytes. Having observed thinning of the corneal stroma in *Zfp469*^BCS/BCS^, we next analysed the amount of Collagen I and Collagen V in the corneal stroma. In wildtype corneas at 6 months of age, immunostaining for both collagen I and V showed uniform distribution throughout the stroma, each having a characteristic appearance (Fig.5A). Type I collagen (ColI) staining appears to demarcate linear structures within the stroma; type V collagen staining (ColV) staining had a more punctate appearance within the linear structures. This is consistent with their roles in assembly of collagen fibrils in the cornea, with collagen type I the predominant fibril-forming collagen, whilst collagen type V has an important role in the nucleation of collagen I fibrils. In *Zfp469*^BCS/BCS^ cornea sections, the staining of collagen I and V was strikingly confined to a much thinner stroma. Both type I and type V collagen staining in homozygous mutant corneas appeared to be more densely packed than in heterozygous or wildtype corneas but quantitation of fluorescent signals determined that total intensity was not significantly different to wildtype controls (data not shown).

**Fig. 5.**
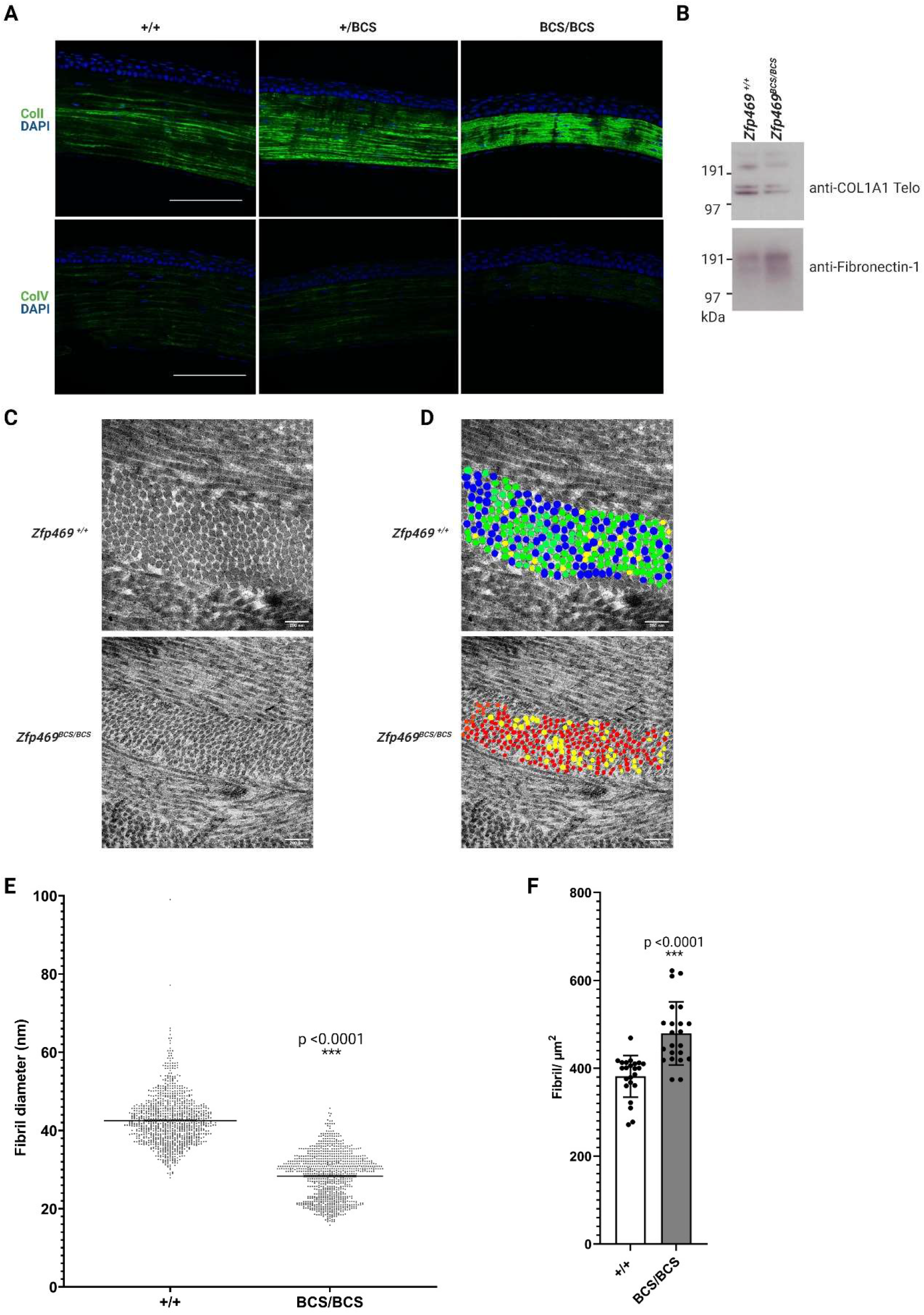
Type I collagen is less abundant and affects fibril architecture in the stroma of *Zfp469*^BCS/BCS^ corneas. (A) Type I and type V collagen localise to the corneal stroma in 6 month old mice, with decreased stromal area apparent in *Zfp469*^BCS/BCS^ compared to wildtype and heterozygous age- and sex-matched samples subject to immunofluorescence microscopy. Nuclei are stained blue with DAPI, collagen type I and type V are stained green. Scale bar is 100 μm. (B) Col1a1 is decreased in *Zfp469*^BCS/BCS^ corneas at 2 months of age by western blot analysis, using fibronectin-1 as a loading control. (C) Representative transmission electron micrographs (TEM) of the cornea anterior stroma show regular fibril organisation and packing of lamellae in mice 9 months of age. Scale bar is 200 nm. (D) Colour coded images of collagen fibrils in the images shown in (C), shows fibrils in wildtype corneal stroma are mostly green (40-49 nm) whereas homozygous mutant fibrils in the anterior stroma are mostly <29 nm (red). Fibrils with diameter 30-39 nm are coded yellow and 50-59 nm are blue. (E) Fibril diameter is significantly decreased in the anterior stroma of *Zfp469*^BCS/BCS^ mice (p,0.0001, Unpaired t test). The range of diameters in 1059 measurements made in sections from 3 wildtype mice was 27.93-99.05 nm, in 912 measurements made in sections from *Zfp469*^BCS/BCS^ mice the range was 15.79-45.70 nm. (F) The smaller fibrils in *Zfp469*^BCS/BCS^ stroma are more tightly packed, as shown by significantly increased fibril density in the corneal stroma of *Zfp469*^BCS/BCS^ mice, measured in 23 sections from 3 wildtype mice and 22 from from *Zfp469*^BCS/BCS^ mice (p,0.0001, Unpaired t test).

Less Col1a1 Telo collagen was detected by immunoblotting in the lysate of 6 pooled *Zfp469*^BCS/BCS^ corneas compared to wildtype, relative to the amount of fibronectin-1 (Fig.5B). These data suggested that collagen production or fibrillogenesis may be affected in BCS corneas, leading us to assess the organisation of the corneal stroma and fibril structure in mice at 9 months of age using transmission electron microscopy (TEM) (Fig.5C). Images obtained from the anterior central cornea showed the lamellae were cylindrical and regularly arranged in both wildtype and mutant corneas, but revealed a striking difference in fibril diameter between 3 wildtype and 3 *Zfp469*^BCS/BCS^ samples. Morphometric analysis revealed that the diameters of cross-sectional collagen fibrils were significantly smaller in *Zfp469*^BCS/BCS^ stroma than in wildtype (mean fibril diameter 28.32 nm ± 5.93 nm s.d. compared with 42.53 nm ± 6.32 nm s.d. respectively, p<0.0001). In anterior stroma of individual wildtype and *Zfp469*^BCS/BCS^ corneas, there is a narrow distribution of fibril size, as shown in the colour coded images in Fig.5D. Distribution of collagen fibril diameter in 3 corneas per genotype group is shown in Fig.5E. Collagen fibril diameter in the anterior stroma was, on average, 33% reduced in *Zfp469*^BCS/BCS^ mice. In addition, an increased number of collagen fibrils were present in a given area of mutant stroma compared to wildtype, with fibril density (fibrils/μm^2^) of the smaller diameter fibrils being significantly increased from 381.7 ± 47.46 (s.d.) fibrils/μm^2^ to 479.3 ± 71.77 (s.d.) fibrils/μm^2^. This corresponds to an increase of 25.6% (Fig. 5F, mutant versus wildtype, p<0.0001) in the stroma of homozygous mutant corneas.

Extraocular features of BCS may include generalised connective tissue dysfunction leading to joint hypermobility, easy bruising and soft, doughy skin (Burkitt Wright et al., 2013). During euthanasia, there were 3 instances of tail degloving (separation of the skin and subcutaneous tissues) in *Zfp469*^BCS/BCS^ adult mice. No such degloving events occurred for wildtype or heterozygous mice. This suggests that LOF of Zfp469 may compromise the biomechanical strength of the skin or underlying connective tissue. Given that type I collagen is the major component of human skin (85–90%), with type III collagen comprising 8-11% and type V collagen 2-5% of total collagen, we sought to determine whether the skin was affected in *Zfp469*^BCS/BCS^ mice. Sections of tail skin from 6 month old mice showed decreased thickness of the dermal and subcutaneous adipose layers in homozygous mutants by H&E staining (Fig.S1A). Masson’s Trichrome staining of collagen in the dermis revealed that the skin of *Zfp469*^BCS/BCS^ mice contains less collagen than wildtype animals (Fig.S1). This appears to arise from decreased abundance of type I collagen in the dermis, as shown in a representative image of adult tail skin in Fig. S1C. In keeping with our earlier observation from corneal sections, type V collagen staining did not alter with genotype (Fig.S1D), suggesting that deficiency of type I collagen underlies both the ocular and extraocular features of BCS.

### A primary cell model of stromal composition in Brittle Cornea Syndrome

Keratocytes synthesise components of the healthy stroma and repair tissue damage. There was no significant difference in the number, or density, of stromal keratocytes in *Zfp469*^BCS/BCS^ corneal sections visualised by DAPI staining of keratocyte nuclei compared to wildtype controls at 6 months of age (Fig. 5A). In order to determine how loss of function of Zfp469 affects the population of stromal keratocytes, we established primary mouse keratocyte cultures from corneas taken from neonatal pups at p2. At this stage in development of the cornea, keratocytes are still dividing and have yet to become quiescent as in mature corneas. From these cultures of corneal stroma fibroblasts, we investigated the gene expression profile in *Zfp469*^BCS/BCS^ primary keratocytes. First, using RT-qPCR we established that the expression of *Zfp469* closely reflected that seen in freshly isolated keratocytes, with no significant difference in *Zfp469* gene expression between genotypes (Fig.6A). Half of the transcript detected in heterozygous cell lines carried the V5 and premature stop CRISPR-Cas9 mediated gene edit; in homozygotes, 100% of the *Zfp469* transcript carried this mutation.

**Fig. 6.**
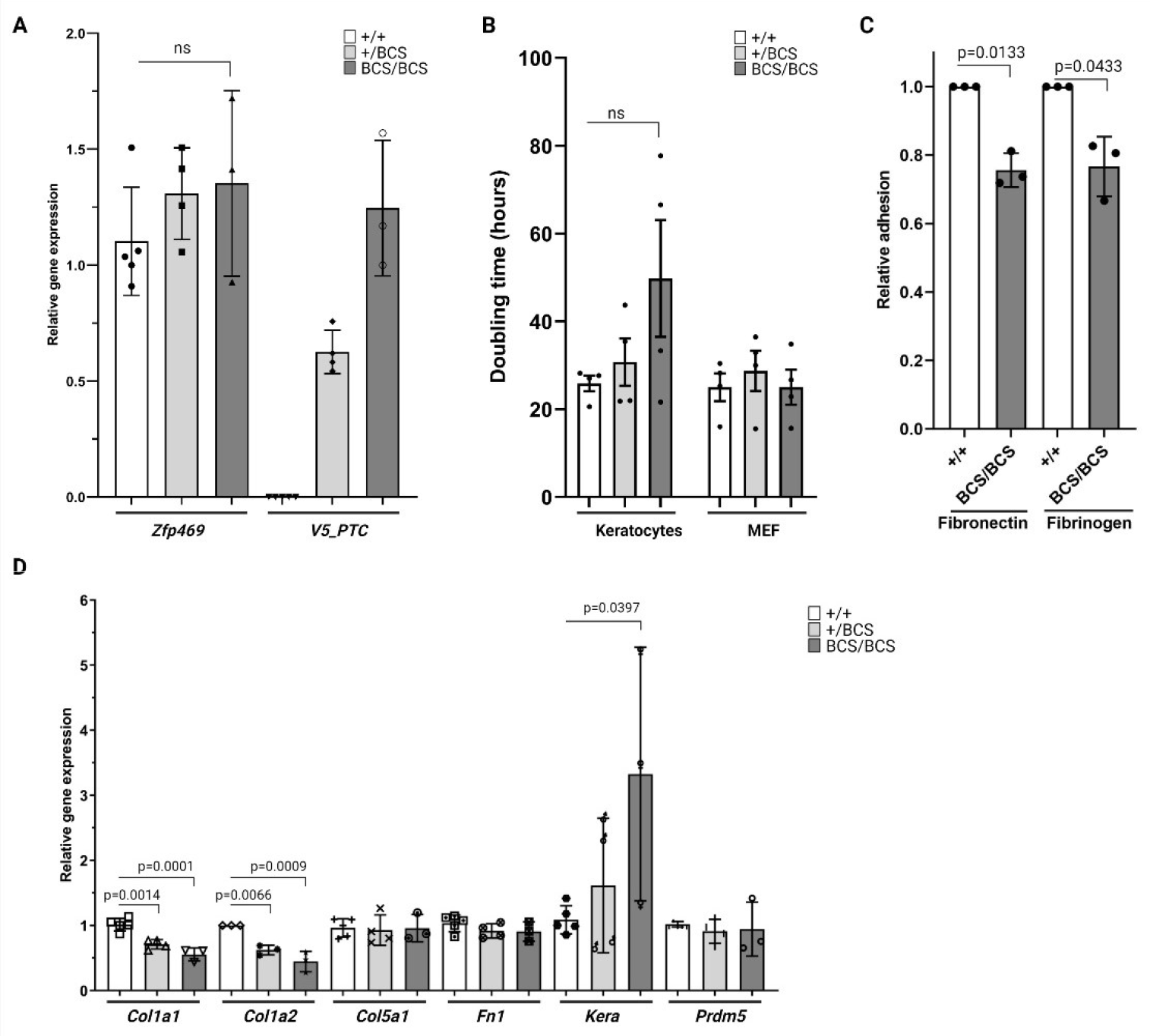
Mutation in Zfp469 affects the expression of ECM genes by corneal stroma fibroblasts in culture. (A) Relative expression of *Zfp469* in primary keratocytes cultures, determined by qRT-PCR, was not significantly different between genotypes. There is an allele dosage effect on the expression of *Zfp469* transcript containing the V5 tag and premature termination codon (V5_PTC), with *Zfp469*^+/BCS^ expressing half as much of this transcript as *Zfp469*^BCS/BCS^ keratocytes lines (n=5 for *Zfp469*^+/+^, n=4 for *Zfp469*^+/BCS^, n=3 for *Zfp469*^BCS/BCS^, one way ANOVA with Dunnett’s multiple comparison test). Data are presented as mean ± s.d., with the average of assays performed in triplicate from each sample shown by data points. (B) Primary keratocytes and MEFs were seeded at known density at passage 3, and the doubling time determined by cell counting after 96 hours in culture. No significant difference in doubling time was observed, although *Zfp469*^BCS/BCS^ keratocytes showed the greatest variability and longest average doubling time. Data are presented as mean ± s.e.m., with the doubling time of individual cell lines shown by data points. (C) Relative adhesion of *Zfp469*^BCS/BCS^ cells to fibronectin and fibrinogen-coated wells after 45 minutes compared to litter-mate wildtype keratocytes was significantly decreased (Wildtype adhesion normalised to 1, average adhesion of *Zfp469*^BCS/BCS^ cells to fibronectin 0.755 ± 0.049 (s.d.) (p=0.0133, Welch’s t-test), average adhesion of *Zfp469*^BCS/BCS^ cells to fibrinogen 0.766 ± 0.087 (s.d.), p=0.0433, Welch’s t-test). Data are presented as relative adhesion (wildtype absorbance 560 nm/Homozygous absorbance 560 nm) ± s.d.. 3 cell lines for each genotype were used. (D) Relative expression of key genes in the ECM and in BCS was assessed by qRT-PCR in primary keratocytes. *Col1a1*, *Col1a2* and *Kera* show significant *Zfp469*^BCS^ allele dosage effect on gene expression. *Hprt* and *Gapdh* were used to normalise the amount of mRNA, and data were analysed using the 2^−ΔΔCT^ method for relative quantitation. Fold change is relative to *Zfp469*^+/+^. Data are presented as mean ± s.d., with the average of assays performed in triplicate from each sample shown by data points (n=5 for *Zfp469*^+/+^, n=4 for *Zfp469*^+/BCS^, n=3 for *Zfp469*^BCS/BCS^, one way ANOVA with Dunnett’s multiple comparison test).

The proliferative ability of the BCS keratocytes *in vitro*, determined using the population doubling time for each cell line seeded at the same density at passage 3, was not significantly altered by the loss-of-function mutation in Zfp469 (Fig. 6B). However, 3 out of 4 *Zfp469*^BCS/BCS^ primary keratocyte lines did proliferate more slowly than wildtype lines isolated from littermates, suggesting that there may be an impact on the rate of proliferation that we are underpowered to detect. The mean doubling time for *Zfp469*^+/+^ primary keratocytes was 25.86 ± 1.77 (s.e.m.) hours compared to 49.80 ± 13.31 (s.e.m.) hours for *Zfp469*^BCS/BCS^ keratocytes (n=4 for each genotype, Fig.6B). Of further note, the 2 mutant cell lines that proliferated most slowly failed to proliferate well beyond passage 4. This was not observed for wildtype cell lines generated from litter mates (data not shown) or for MEFs, which showed no variability in doubling time by genotype (Fig. 6B). This suggests that there might be a genotype, and cell-type, specific effect of *Zfp469* mutation on cell proliferation.

The adhesion of keratocytes to ECM is key to establishing and maintaining the structural integrity of the cornea (Parapuram et al., 2011). We sought to determine the impact of LOF of Zpf469 on adhesion of primary keratocytes to a variety of ECM components, including Collagen I, Collagen IV, Fibrinogen, Fibronectin and Laminin. This revealed that *Zfp469*^BCS/BCS^ keratocytes adhere less efficiently to fibrinogen (p = 0.0433) and fibronectin (p = 0.0133) compared to wildtype controls (Fig.6C). Defective adhesion to fibronectin, an ECM protein with known roles in maintenance of tissue architecture and wound healing, may impact upon arrangement and organisation of collagen fibrils in the stroma.

The expression of *Prdm5*, in which mutations cause BCS type 2, and key components of the stromal ECM was investigated by qPCR (Fig. 6D). *Prdm5* expression was unchanged in *Zfp469*^BCS^ primary keratocytes. However, a significant impact of *Zfp469*^BCS^ allele dosage on the expression of *Col1a1* and *Col1a2*, the genes encoding the alpha 1 and alpha 2 chains of collagen type I that associate to form the triple helix of individual type I collagen, was uncovered. The relative expression of both *Col1a1* and *Col1a2* was reduced by approximately 50% in homozygous mutant cell lines compared to wildtype lines (*Col1a1*, adjusted p=0.0001; *Col1a2*, adjusted p=0.0009, one way ANOVA with Dunnett’s multiple comparison test), and by approximately 25% in heterozygous primary keratocytes (*Col1a1*, adjusted p=0.0014; *Col1a2*, adjusted p=0.0066) (Fig.6D). The expression of *Col5a1* in BCS keratocytes in culture was not significantly altered compared to wildtype cell lines, agreeing with our observations from staining of corneal sections. Expression of *Kera*, a key keratocyte marker, also showed an allele dosage effect of mutation of Zfp469, increasing 1.5 fold in ^+/BCS^ and 3-fold in ^BCS/BCS^ keratocyte cell lines (adjusted p=0.0397, one way ANOVA with Dunnett’s multiple comparison test). *Kera* encodes keratocan, a cornea-specific keratan sulfate proteoglycan (KSPG) belonging to the small leucine-rich proteoglycan (SLRP) gene family and it has an important role in regulation of collagen fibril spacing and arrangement in the mature cornea. These data point to a dysfunctional gene expression profile in mutant keratocyte cells that may affect the composition and assembly of the stromal extracellular matrix.

We next investigated the secretion and deposition of type I collagen by primary keratocytes. The amount of proCol1a1 secreted into the medium by confluent *Zfp469*^BCS/BCS^ primary keratocytes maintained in serum-free medium for 7 days was significantly reduced compared to wildtype cells (Fig.7A, p=0.0332, t-test). Immunoblotting of the same serum-free conditioned medium samples using an anti-Telo Col1a1 antibody confirmed that *Zfp469*^BCS/BCS^ primary keratocytes secrete less Col1a1 (Fig.7B) with densitometry revealing a 43 ± 9 (s.d.) % depletion of Col1a1 secreted relative to wildtype samples. Cell-derived matrices (CDM) from homozygous mutant and wildtype primary keratocytes were generated to study ECM deposition *in vitro*. Keratocytes were seeded onto gelatin-coated plates or coverslips and, once confluent, were treated with ascorbic acid for 20 days in order to induce deposition of a collagen-rich matrix. Immunoblot analysis of the CDMs collected following decellularisation under reducing and denaturing conditions revealed that less type I collagen was deposited by mutant cell lines (Fig. 7C). The expression of the fibronectin-1 gene (*Fn1*) was unaffected by Zfp469 mutation (Fig. 6D). Using densitometric analysis of Col1a1 Telo signal normalised to fibronectin-1 (Fig. 7D), *Zfp469*^BCS/BCS^ primary keratocytes deposited, on average, only 30% of Col1a1 to the CDM relative to wildtype cells (p=0.0097, t-test). Intact, decellularised CDMs were also stained with antibodies against Collagen type I, Collagen type V and Fibronectin (Fig. 7E). Quantification of fluorescent signals in CDM generated by two *Zfp469*^BCS/BCS^ cell lines confirmed that less type I collagen was deposited by *Zfp469*^BCS/BCS^ primary keratocytes, whilst collagen type V remained unchanged relative to wildtype (Fig. 7F).

**Fig. 7.**
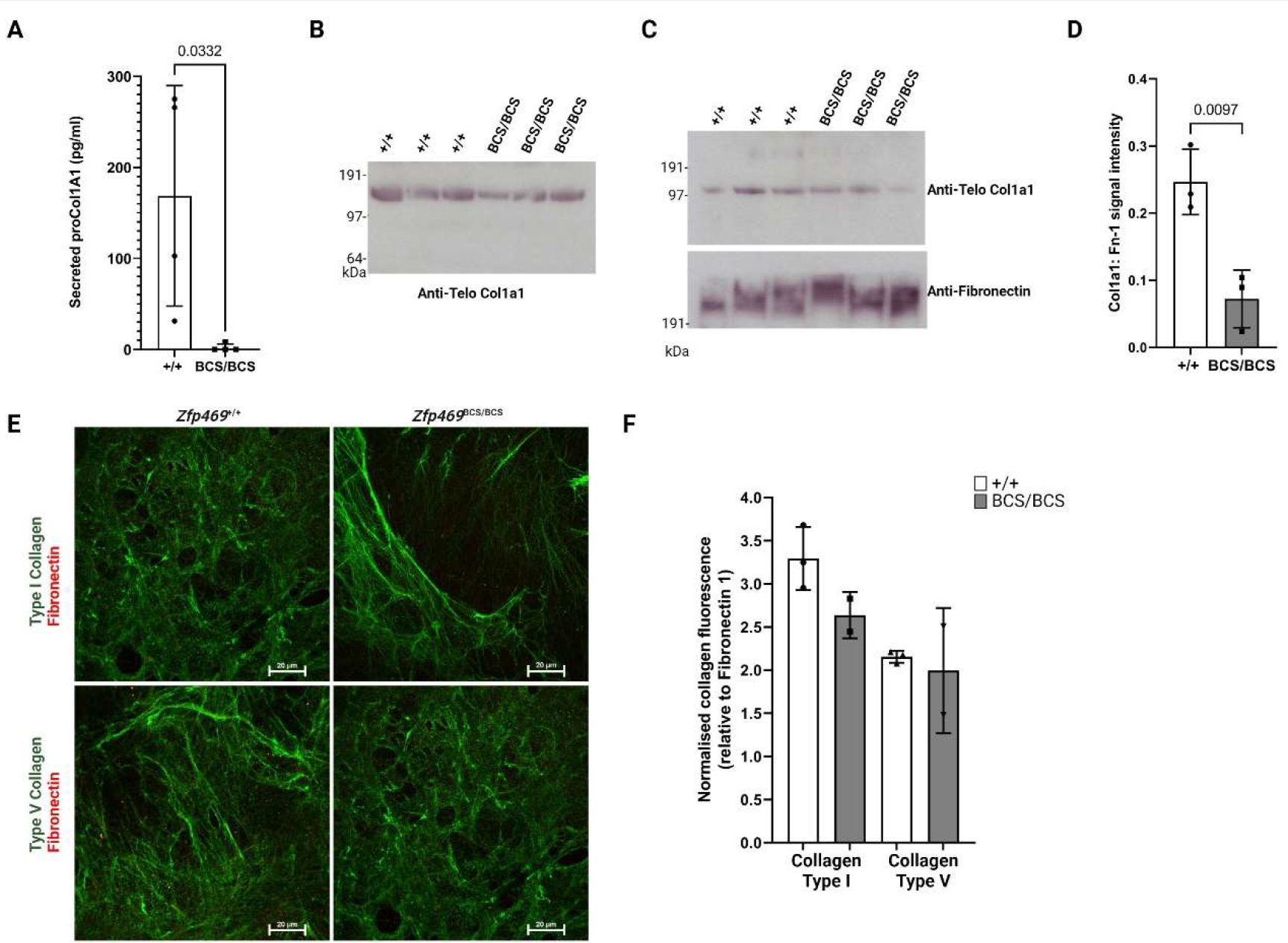
Secretion and deposition of type I collagen by primary keratocytes is impaired when Zfp469 is mutated. (A) The concentration of proCol1a1 in serum-free media after 7 days in culture was measured by ELISA showing that secretion of proCol1a1 into culture medium by confluent *Zfp469*^BCS/BCS^ primary keratocytes was significant impaired compared to wildtype cells. Data are presented as mean ± s.d. with the proCol1a1 concentration of individual cell lines shown by data points (n=4 for *Zfp469*^+/+^, n=4 for *Zfp469*^BCS/BCS^, t-test). (B) Representative western blot of the serum-free conditioned medium samples used in ELISA probed with an anti-Telo Col1a1 antibody which detects prepro-, pro- and mature Col1a1 shows decreased Col1a1 in the medium of *Zfp469*^BCS/BCS^. (C) Cell-derived matrices (CDM) generated by primary keratocytes were decellularised and then denatured and reduced prior to western blotting. The CDM of *Zfp469*^BCS/BCS^ contained less Telo Col1a1 than wildtype CDM, using fibronectin as a loading control. (D) Quantification of Col1a1 signal normalised to fibronectin for CDMs generated from 3 *Zfp469*^+/+^ and 3 *Zfp469*^BCS/BCS^ by densitometry shows a significant decrease in abundance of Telo Col1a1 in mutant cell lines (n=3 per group, p= 0.0097, unpaired two-tailed t test). (E) Representative immunofluorescence images of CDMs generated by primary keratocytes after 20 days exposure to ascorbic acid-containing medium, stained for type I collagen (green), type V collagen (green) and fibronectin (red). Scale bar represents 20 μm. (F) Immunofluorescence staining intensity for type I collagen and type V collagen was normalised to fibronectin signal for CDMs from 3 *Zfp469*^+/+^ and 2 *Zfp469*^BCS/BCS^ cell lines, using 5 non-overlapping images from each sample. Data are presented as mean ± s.d. with the average normalised collagen staining intensity of individual cell lines shown by data points. Deposition of type V collagen did not differ between genotypes, but type I collagen deposition was decreased in *Zfp469*^BCS/BCS^ CDMs.

## Discussion

Brittle cornea syndrome was first reported in 1968 (Stein et al., 1968) but it was another 40 years until mutations in *ZNF469* were identified as a cause of this rare condition. Since then, 30 pathogenic mutations in *ZNF469* have been identified, the majority of these resulting in premature stop codons preceding ZF domains. The precise mechanism by which biallelic loss-of-function mutations result in generalised connective tissue dysfunction, and why the most severe manifestation occurs in the cornea and sclera, has remained unclear. This study is the first description of a mouse model of BCS, in which we created a premature stop codon in the orthologous mouse gene *Zfp469* to recapitulate the human mutation p.Gly677*. Our results showed that this premature stop codon in the large single coding exon of *Zfp469* does not lead to nonsense mediated decay of the mutant transcript and will likely lead to generation of a truncated protein. Despite incorporation of an in-frame V5 tag, we were unable to detect a V5-tagged protein by western blot or immunostaining of tissue samples or fixed primary keratocytes. This suggests that the protein may be expressed at very low levels, the mutated V5 is not recognised or that the V5 tag is inaccessible. Although the mutant protein could not be detected, disrupting Zfp469 clearly perturbs the development of the corneal stroma in mouse, resulting in thinning of the cornea by one third in homozygous mutant eyes at 1 month of age. At 3 and 6 months of age, the mature cornea of *Zfp469*^BCS/BCS^ eyes remain 30-35% thinner than wildtypes, suggesting that, in mice, stromal thinning is not progressive.

BCS patients suffer from keratoconus, keratoglobus, corneal edema resulting from breaks in the Descemet’s membrane, and extremely thin corneas are prone to rupture after minor injury (Burkitt Wright et al., 2013). Although none of our BCS mice, aged up to 9 months, suffered from cornea rupture, one case of corneal edema was recorded. The bulging eye phenotype observed in our BCS mouse model following dilation of the pupil closely resembled keratoglobus. Furthermore, the corneas of homozygous mutant mice showed a decreased ability to withstand corneal strain, deforming after either application of an external eye drop or by dilation of the pupil. This is consistent with a loss of biomechanical strength in the corneal stroma when Zfp469 function is lost, and is in striking contrast to the corneas of wildtype mice, which were able to maintain normal corneal curvature under the same pressure load. This study is the first to show a direct impact of LOF of Zfp469 upon cornea biomechanical strength. However, the genetic association of regulatory variants for *ZNF469* with corneal resistance factor (CRF) in genome-wide association studies indicates that ZNF469 also plays a role in determining biomechanical properties of the cornea in humans (Jiang et al., 2020; Khawaja et al., 2019; Simcoe et al., 2020).

Numerous studies have shown that disrupting the composition or organisation of the stromal extracellular matrix will affect the shape (Quantock et al., 2003) or strength (Chakravarti et al., 1998) of the cornea. Consistent with this, the corneal thinning caused by loss of Zfp469 appears to result from approximately 50% reduction in the expression of genes encoding type I collagen, *Col1a1* and *Col1a2*, by keratocytes. This is an important result, as the relative abundance of type I and type V collagen in the cornea appears key to the proper organisation of the stroma. Collagen composition in the cornea is unique amongst tissues, with fibrillar collagen in the stroma made up of approximately 80-90% type I and 10-20% type V. Type I collagen assembles into heterotrimers of two alpha 1(I) chains and one alpha 2(I) chain, growing further into heterotypic fibrils with type V collagen. Non-helical terminal extensions of type V collagen protrude from the fibril (Linsenmayer et al., 1993) acting to nucleate fibril growth and determine fibril diameter. Strikingly, the expression of collagen type V remained unchanged in our primary keratocytes cells, in cell derived matrices, and by staining of corneal sections from *Zfp469*^+/+^ and *Zfp469*^BCS/BCS^ mice, providing possible insight into the pathomechanisms underlying BCS caused by mutation in Zfp469.

Analysis of fibril diameter, density and organisation in the corneal stroma of our BCS model by TEM further implicated reduced type I collagen expression in corneal stromal thinning. The diameter of collagen fibrils was reduced by 33% in *Zfp469*^BCS/BCS^ mice compared to wildtype and fibril density was increased by 25%. This closely resembled the ultrastructural phenotype of the Osteogenesis Imperfecta (OI) mouse model resulting from homozygous mutation in *Col1a2* (*oim*, (Chipman et al., 1993)). CCT of *Col1a2^oim/oim^* corneas is 15% thinner than wildtype controls, and fibril diameter determined by TEM is also reduced by 15%. These features arise from impaired collagen fibrillogenesis as a result of loss of Col1a2 during the assembly of type I collagen heterotrimers (Dimasi et al., 2010). Notably, type I collagen homotrimers containing 3 units of Col1a1 have been observed in adult skin and in OI (Nicholls et al., 1984; Pace et al., 2008; Pihlajaniemi et al., 1984; Uitto, 1979). The concurrent decrease in expression of both *Col1a1* and *Col1a2* may explain the more severe corneal thinning observed in both the BCS model and in BCS compared to OI in humans.

Interestingly, arrangement of fibrils within the lamellae of mature *Zfp469*^BCS/BCS^ corneas was regular and the organisation of parallel lamellae appeared normal. Indeed, corneal transparency was maintained throughout our study. A similar observation has been made in a previous report of *Col5a1* haploinsufficiency, where heterozygous loss of *Col5a1* results in thinning of the mature mouse corneal stroma by 26%, resulting in 14% decrease in total collagen content and a 25% reduction in the number of fibrils in the stroma. Fibril diameter was increased and fibril density was reduced in this model of Ehlers-Danlos Syndrome, but no corneal opacity was observed (Segev et al., 2006). This is in contrast to Col5a1^Δst/Δst^ mice, where targeted deletion of *Col5a1* in the corneal stroma results in total absence of Col5a1 from the stroma, leading to larger and less uniform collagen fibrils and extensive corneal hazing (Sun et al., 2011). These data confirm, firstly, that regular packing of a homogeneous population of fibrils with a narrow distribution of diameters is key to transparency (Hassell and Birk, 2010), and secondly, that the ratio between type I and type V collagen is crucial to the assembly of a well-ordered and mechanically strong corneal stroma.

Despite the evident allele dosage effect upon the expression of both *Col1a1* and *Col1a2* by keratocytes, with heterozygous cell lines expressing an intermediate amount relative to wildtype and homozygous lines, the corneas of heterozygous *Zfp469*^BCS^ mice are only slightly thinner than wildtype. This mimics the observations from heterozygous carriers of LOF *ZNF469* and *PRDM5* mutations, who appear asymptomatic (Abu et al., 2008; Alazami et al., 2016; Aldahmesh et al., 2012; Avgitidou et al., 2015; Burkitt Wright et al., 2011; Christensen et al., 2010; Khan et al., 2012; Menzel-Severing et al., 2019; Micheal et al., 2016; Ramappa et al., 2014; Rohrbach et al., 2013; Skalicka et al., 2020) or show only mild corneal thinning and blue sclera (Burkitt Wright et al., 2011; Khan et al., 2010). This may reflect a critical threshold for the relative abundance of type V collagen to type I collagen in the cornea, which is known to have a major role in determining both the number of fibrils that form and determining the diameter of fibrils (Dimasi et al., 2010; Sun et al., 2011; Wenstrup et al., 2004). Compared to skin, where type V collagen comprises only 2-5% of total fibrillar collagen, the relative ratio of type V to type I collagen in the cornea is normally 5-10 fold higher. In our mouse model of BCS, this relative ratio is increased even further as the amount of type I collagen is reduced in both the corneal stroma and in skin. *ZNF469* mutations have been shown to disrupt deposition of type I collagen by dermal fibroblasts (Burkitt Wright et al., 2011), in agreement with our observation of decreased dermal thickness and type I collagen abundance in tail skin from *Zfp469*^BCS/BCS^ mice. However, this is the first report of mutation in Zfp469 disrupting the expression of type I collagen in the corneal stroma, and by primary keratocytes. Consistent with the role of collagen type V as a rate-limiting fibril nucleator, this appears to promote fibril nucleation from the depleted pool of type I collagen synthesised by keratocytes in *Zfp469*^BCS/BCS^ corneal stroma. This results in the extremely thin corneal stroma, comprising smaller diameter and more tightly packed collagen fibrils than in wildtype mice, explaining the phenotype observed in our model of the disease and in BCS patients.

Extreme thinning of the cornea was observed at 1 month of age in *Zfp469*^BCS/BCS^ mice, and BCS cases have been reported in humans soon after birth (Ramappa et al., 2014) suggesting that this disorder is initiated during development. This has also been observed in a mouse model for Marfan syndrome, another connective tissue disorder, which shows reduced CCT in *Fbn1^+/−^* mice from E16.5 onwards (Feneck et al., 2020). The corneal stroma is established during early development, with collagen deposition by presumptive corneal stromal cells observed at E13 in the mouse (Feneck et al., 2019). As development proceeds, organised collagen fibrils begin to form, directed by keratocyte cell extensions into parallel arrangement. Type I collagen appears to play a key role in this process, as complete knockout of *Col1a1* in the Mov13 mouse shows reduced corneal thickness at E16 and structural disorganisation of developing fibrils (Bard and Kratochwil, 1987). As the cornea matures, up to 8 weeks of age in the mouse, or 3 years of age in humans, collagen fibrils are further organised into additional layers of the established lamella. Our work suggests that the corneal thinning observed in *Zfp469*^BCS/BCS^ mice arises from a deficiency of type I collagen expression by keratocytes due to mutation in *Zfp469*. There does not appear to be a paucity of keratocytes in the stroma of mature cornea, but we cannot exclude an impact on differentiation from neural crest cells or upon proliferation of keratocytes at early stages of development. Future work using our mouse and cellular models of disease will determine whether corneal thinning in BCS is established during very early stages of development, how ZNF469 regulates expression of type I collagen genes, and if there is a therapeutic window of opportunity for the modulation of ZNF469 function.

## Materials and Methods

### Bioinformatics

A literature review was performed to identify mutations in ZNF469 that have been reported in BCS cases. Using the search term “ZNF469”, 12 case reports reporting ZNF469 mutations were identified, with a total of 30 different mutations occurring either as compound heterozygotes or homozygous in 53 cases of BCS were implicated. All mutations were mapped to transcript NM_001367624.2 and protein NP_001354553.1 using Ensembl Variant Effect Predictor (VEP, https://www.ensembl.org/Tools/VEP). The Genome Aggregation Database (gnomAD v3.1; https://gnomad.broadinstitute.org/) was searched for BCS mutations that may be present in the general population. Protein sequence alignments were performed using Clustal Omega (https://www.ebi.ac.uk/Tools/msa/clustalo/). Uniprot (https://www.uniprot.org/) and SMART (http://smart.embl-heidelberg.de/) were used to identify positions of C2H2 zinc finger domains and compositional bias in the amino acid sequence.

### Preparation of reagents for genome editing

Paired sgRNA for use with nickase Cas9 were designed using the tool available at http://crispr.mit.edu (May 2017). Three pairs were designed and top and bottom strands ordered as 5’ Phosphorylated single stranded DNA oligos (IDT) with BbsI-cloning compatible ends for cloning into pX458 as previously described(Ran et al., 2013). pSpCas9(BB)-2A-GFP (PX458) was a gift from Feng Zhang (Addgene plasmid # 48138 ; http://n2t.net/addgene:48138 ; RRID:Addgene_48138). Sequences of sgRNA are shown in Table S1. A single-stranded donor oligonucleotide (ssODN) repair template with 5’-56 base pair and 3’-54 base pair homology arms was designed to insert a V5 epitope tag and a premature stop codon into mouse Zfp469 at p.Gly634, and ordered as a PAGE Ultramer DNA Oligo (IDT). After verification of correct sequence by Sanger sequencing, plasmid DNA for each sgRNA was used to test efficiency of on-target editing at the *Zfp469* locus in mouse embryonic fibroblasts (MEFs). After selection of the most efficient pair of sgRNA, plasmid DNA was used as a template in PCR to amplify sgRNA 1 and 3 sequences using primers shown in Table S1. DNA was purified using Qiagen PCR purification columns, and 1 μg of purified PCR product was used as template for *in vitro* transcription using the HiScribe T7 High Yield RNA Synthesis kit (New England Biolabs, E2040S) as described by the manufacturer.

### Testing sgRNA in mouse embryonic fibroblasts

Wildtype MEFs (200,000/reaction) were transfected with 2 μg pX458 plasmid containing sgRNA using Neon electroporation (1350V, 30 ms and 1 Pulse). Transfected cells were incubated for 72 hours in a humidified 37°C, 5% CO_2_ incubator after which cells were collected. The efficiency of sgRNA-mediated DNA cleavage at the on-target locus was determined using a GeneArt Genomic Cleavage Detection Kit (Invitrogen) as described by the manufacturer. Primers used for target amplicon amplification were Forward 5’-TTCATCTCTGTCACCGCCAT-3’ and Reverse 5’-GAAGGGGACAGTCTGGTTGT-3’.

### Genome editing in mice

Cytoplasmic injection of 20 μl containing 50 ng/μl SpCas9n mRNA, 50 ng/μl sgRNA, and 150 ng/μl donor oligonucleotides in to C57BL/6J mouse zygotes was performed, followed by immediate transfer to pseudopregnant CD1 females. Founder mice were genotyped by extracting DNA from earclips in DNAreleasy (Anachem) following the manufacturer’s instructions. Primers (Forward 5’-TTCATCTCTGTCACCGCCAT-3’ and Reverse 5’-GAAGGGGACAGTCTGGTTGT-3’) were used to amplify DNA surrounding the editing target site from crude DNA by PCR using DreamTaq Green PCR Master Mix (Thermo Scientific). Sanger sequencing was used to verify editing and repair. One founder mouse with in-frame repair was identified and was used to establish the *Zfp469*^BCS^ line by breeding with wildtype (WT, ^+/+^) C57BL/6J.

### Mouse husbandry

Experiments on mice were performed with UK Home Office project licence approval. Animals were housed in a facility on a 12 hour light/dark cycle with unrestricted access to food and water. All mice were euthanised in accordance to UK Home Office guidelines. Heterozygous F1 offspring of the founder mouse were interbred to generate subsequent generations of *Zfp469*^BCS^ mice used in this study, maintained on the C57BL/6J strain background. Litters were genotyped as for the founder mice, or outsourced to Transnetyx (Cordova, TN, USA) using allele-specific custom probes. Male and female mice were used in this study.

### Slit lamp examination

The anterior segment of the eyes of 3 *Zfp469*^+/+^, 3 *Zfp469^+/^*^BCS^ and 3 *Zfp469*^BCS/BCS^ mice were examined at 3 months of age using a slit lamp biomicroscope. Mice were examined without anaesthesia, and images taken with a digital camera.

### Measurement of eye size

Freshly enucleated eyes from 2 homozygous mice and 3 wildtype mice were placed upon a flat surface, and measured from the front of the cornea to the posterior just left of the nerve four times using calipers. An average measurement was calculated for each eye.

### Optical Coherence Tomography

Anterior segment OCT images were captured using a Heidelberg Spectralis OCT. Mice aged 1 month, 3 months and 6 months of age were anaesthetized by inhalation of isoflurane, and corneas were imaged using an anterior segment lens. To test corneal distortion after pupil dilation, a drop of Tropicamide 1% and then a drop of 2.5% Phenylephrine Hydrochloride was added to each eye prior to AS-OCT imaging. Systane Ultra Lubricant (Boots) eye drops were used to prevent eyes drying out.

Corneal dimensions including central and peripheral corneal thickness and anterior chamber depth were determined from cross-sectional corneal images that passed through the centre of the pupil using ImageJ software (National Institutes of Health). CCT and PCT were determined by measuring the linear distance between the anterior and posterior corneal surfaces in the central cornea and peripheral corneal respectively. ACD was the distance between the lens and the posterior surface of the central cornea. The measurements were obtained in pixels and the appropriate pixel to μm conversion factor was applied, relative to 200 μm scale bar. One measurement was made for each scan of both eyes for each mouse. A minimum of 3 male and 3 female mice per genotype were used at each 1, 3 and 6 months of age.

### Transmission Electron Microscopy

For Transmission Electron Microscopy (TEM), corneas from three *Zfp469*^+/+^ and 3 *Zfp469*^BCS/BCS^ female mice aged 9 months were removed and fixed in 3% glutaraldehyde in 0.1M Sodium Cacodylate buffer, pH 7.3, for 2 hours then washed in three 10 minute changes of 0.1M Sodium Cacodylate. Corneas were post-fixed in 1% Osmium Tetroxide in 0.1M Sodium Cacodylate for 45 minutes, and washed in three 10 minute changes of 0.1M Sodium Cacodylate buffer. These samples were dehydrated in 50%, 70%, 90% and 100% ethanol (×3) for 15 minutes each, then in two 10-minute changes in Propylene Oxide. Samples were subsequently embedded in TAAB 812 resin. Sections, 1 μm thick, were cut on a Leica Ultracut ultramicrotome, stained with Toluidine Blue, and viewed in a light microscope to select suitable areas for investigation. Ultrathin sections, 60 nm thick, were cut from selected areas, stained in Uranyl Acetate and Lead Citrate and then viewed in a JEOL JEM-1400 Plus TEM. Representative images were collected on a GATAN OneView camera. Multiple high-magnification (× 35,000), non-overlapping cross-sectional images were taken from the anterior stroma in the central cornea of each sample. Fibril dimensions were determined using ImageJ software (National Institutes of Health) to analyse multiple images per sample across a defined region of interest in each image. Fibril diameter was measured by taking two measures perpendicular to each other and an average used for each fibril. Fibril density was calculated as the number of fibrils in the known area of the region of interest in each image. Microsoft Excel and GraphPad PRISM were used for data analysis. Imaging and measurements were performed blinded to genotype.

### Histology

Mice were euthanised at the appropriate age by cervical dislocation before eyes were enucleated and tails were removed for fixing in in 4% Paraformaldehyde (PFA) solution overnight at 4°C. Following this, tails were subjected to 3 washes of 6% EDTA for one week each. For wax preservation, tissue samples were removed from PFA or EDTA, and were subsequently dehydrated by successive washes in 70%, 80%, 90% and 100% ethanol, xylene twice and then embedded in paraffin for 45 minutes.

Hematoxylin and Eosin (H&E) and Masson’s Trichrome staining were performed on 8 μm paraffin embedded tissue sections using standard procedures. Images were captured on a Zeiss Axioplan 2 brightfield microscope. Immunostaining was performed using 5-8 μm paraffin-embedded sections after antigen retrieval in Citrate Buffer, pH6. Slides were rinsed with water, washed twice with Tris-HCl buffered saline (TBS) with 0.1% Tween-20 in (TBST) and then blocked in 4% heat-inactivated donkey serum (DS) in TBST for 30 minutes at room temperature. Primary antibodies (Goat Anti-Type I Collagen, 1310-01, Southern Biotech; Goat Anti-Type V Collagen, 1350-01, Southern Biotech) were diluted 1:100 in 4% DS in TBST and incubated overnight at 4°C. After 3 washes in TBST, Alexa Fluor secondary antibody diluted 1:250 4% DS in TBST was added to the slides for 1.5 hours at room temperature. Slides were washed 3 times in Phosphate Buffered Saline (PBS) prior to mounting coverslips using Prolong Gold Antifade Mountant with DAPI (Invitrogen). Slides were imaged on a Nikon Confocal A1R microscope for image processing using NIS-Elements or ImageJ software.

### Corneal protein extraction

For proteomic analysis, 6 pooled corneas (from different 3 animals at 2 months of age) for each genotype were subject to mechanical grinding and 5 rounds of freeze-thawing in RIPA buffer (150 mM NaCl, 50 mM Tris pH 8, 1% NP-40, 0.5% sodium deoxycholate, 0.1% SDS) plus Complete EDTA-free protease inhibitor cocktail (Roche). After centrifugation at 13,000 rpm for 10 minutes at 4°C, supernatants were recovered and protein concentration determined using BioRad Protein assay according to the manufacturer’s instructions, with BSA as standard.

### Gel electrophoresis and western blotting

Samples containing equal amounts of total protein were mixed with 1 × Reducing Agent (Invitrogen by ThermoFisher) and 4 × NuPAGE LDS (lithium dodecyl sulfate) Sample Buffer (Invitrogen by ThermoFisher). Samples were denatured by heating to 80°C for 15 minutes. Gel electrophoresis was performed using 4–12% Bis-Tris NuPAGE gels (Invitrogen by ThermoFisher) run in 1 × MOPS (3-(N-morpholino)propanesulfonic acid; Invitrogen by ThermoFisher). Proteins were transferred to Hybond-P PVDF membranes (GE Healthcare) in 1 × NuPAGE Transfer Buffer (Invitrogen by ThermoFisher) containing 20% methanol (Fisher). Membranes were blocked in 5% non-fat milk in PBST (Phosphate-buffered Saline-Tween 20; 3.2 mM Na_2_HPO_4_, 0.5 mM KH_2_PO_4_, 1.3 mM KCl, 135 mM NaCl, 0.05% Tween 20, pH 7.4) before being incubated with primary antibodies (Anti-Telo-Collagen Type I, A1, ABT256, Sigma-Aldrich; Anti-Fibronectin-1, F3648, Sigma-Aldrich) followed by anti-rabbit IgG horseradish peroxidase (HRP)-conjugated secondary antibody. Protein bands were visualised using Enhanced Chemiluminescence (ECL) Western blotting detection reagents (GE Healthcare) and exposure of the membrane to Kodak BioMax XAR film (Sigma). Films were developed using a phosphorimager X-ray machine (Konica Minolta). Densitometric analysis of scanned films was performed using ImageStudioLite software (Li-Cor).

### Mouse embryonic fibroblast generation and culture

MEFs were isolated from E13.5 embryos from heterozygous *Zfp469^+/^*^BCS^ crosses. Heads and organs were removed from embryos and the remaining sample was digested with trypsin for 10 minutes at 37 °C to generate a cell suspension. Cells were pelleted by centrifugation at 1200 rpm for 4 minutes and then cultured in OPTIMEM + 10% FCS + Pen/Strep. MEFs were used between passages 3-6.

### Mouse primary keratocyte generation and culture

Mouse corneal stroma keratocytes were isolated as previously described (Zhang et al., 2016), with slight modification. In brief, pups were euthanised at postnatal day 2, eyes were dissected, rinsed in PBS and corneas cut out along scleral rim. Two corneas (per mouse) were incubated in 15 mg/ml Dispase (4942078001, Roche) in DMEM (Gibco) +Penicillin/Streptomycin for 30 minutes at room temperature. Epithelial cells were detached by gentle shaking and the corneas rinsed in PBS, before incubation in Digestion Buffer containing 2 mg/ml Collagenase (Gibco) and 0.5 mg/ml hyaluronidase (Sigma-Aldrich) in DMEM at 37 °C for 30 minutes, with vortexing every 5 minutes. Samples were centrifuged at 300 × g for 5 minutes before the supernatant was removed, and trypsin/versene added for a further incubation at 37 °C for 10 minutes. Cells were recovered by centrifugation at 300 × g for 5 minutes, supernatant discarded and cells resuspended in culture medium (DMEM, 10% Fetal Calf Serum (FCS), 1% Pen/Strep) for plating. Primary keratocytes (corneal stromal fibroblasts) were maintained in a humidified 37°C, 5% CO_2_ incubator, sub-cultured at 80% confluency, and were used between passage 3-6 during this study.

### Isolation of RNA, cDNA synthesis and real-time quantitative PCR (qPCR)

RNA was isolated from corneal samples dissected from 4 wildtype, 2 heterozygous and 4 homozygous mice aged 3 months and the corneas processed as described to obtain cells for primary keratocytes culture. After treatment with trypsin/versene, the resulting cell pellet was resuspended in Buffer RLT and total RNA prepared using an RNEasy Kit (Qiagen). RNA was also purified from primary mouse keratocytes isolated from p2 pups at passage 4 using the RNEasy kit. Reverse transcription was performed using the Transcriptor High Fidelity cDNA synthesis kit (Roche) and random hexamer primer according to manufacturer’s instructions. qPCR was performed using Taqman Gene Expression assays (*Hprt*, Mm01324427_M1, *Col1a1*, Mm00801666_g1, *Col1a2*, Mm00483888_m1, Col5a1, Mm00489229_m1; *Kera*, Mm00515230; *Fn1*, Mm01256744_m1; *Prdm5*, Mm00510567_m1; *Gapdh,* Mm99999915_g1; custom assays targeting start of the *Zfp469* transcript, and the V5 insertion; Applied Biosystems) with Taqman Universal MasterMix II, no UNG (Applied Biosystems). Assays were run on an ABI PRISM 7900 thermocycler with each sample in triplicate. *Hprt* and *Gapdh* were used as housekeeping genes, and data were analysed using the 2^−ΔΔCT^ method for relative quantitation.

### Proliferation assays

Proliferation assays were performed using 50,000 primary mouse keratocytes or MEF from four independent cell lines for each genotype at passage 3. After 96 hours in culture complete culture medium the cells were trypsinised and counted using a haemocytometer. The number of population doublings in 96 hours was used to calculate the doubling time for each cell line in hours.

### Adhesion assay

The adhesion of primary mouse keratocytes to the ECM components Collagen I, Collagen IV, Fibrinogen, Fibronectin and Laminin was quantitatively tested using a CytoSelect Adhesion Assay Kit (Cell Biolabs, San Diego, CA) as described by the manufacturer. In brief, three wildtype lines, obtained from three different mice, were compared to three homozygous mutant lines, also obtained from three different mice. 75,000 cells in serum-free DMEM were seeded into wells of a 48-well plate pre-coated with the ECM components and incubated for 45 minutes at 37°C. Media was removed and the plate washed 5 times with PBS to remove non-adherent cells, before addition of Cell Stain Solution to each well. After incubation for 10 minutes at room temperature, wells were washed 5 times with PBS and allowed to air dry. Extraction Solution was added and after 10 minutes incubation with shaking, 150 μl of each sample was used to measure absorbance at 560 nm in a TECAN M200 Pro plate reader (TECAN, Switzerland).

### Pro-Collagen 1A1 Enzyme-linked Immunosorbent Assay (ELISA)

Primary mouse keratocytes were seeded 75,000 cells per 10 cm plate and grown to confluency in medium containing FCS. Cells were washed in PBS, and medium changed to serum-free for a further 7 days. Cell culture supernatants were collected and assayed for the secretion of pro-Col1A1 using Human Pro-Collagen I alpha 1 DuoSet ELISA kit (DY6220-05, R&D Systems) as described by the manufacturer.

### Generation of Cell-Derived Matrices (CDM)

Primary mouse keratocytes cells were used to generate CDMs following a published protocol (Kaukonen et al., 2017). In brief, multi-well tissue culture plates and glass coverslips were coated with sterile 0.2% (w/v) gelatin diluted in PBS for 1 hour at 37°C. Coverslips were washed with PBS and then cross-linked with 1% (v/v) glutaraldehyde for 30 minutes at room temperature, prior to quenching with sterile 1 M glycine for 20 minutes. Coverslips and plates were then washed with PBS and incubated in DMEM, 10% FCS, 1% Pen/Strep for 1 hour before use.

Primary mouse keratocytes were seeded 2×10^5^ cells/well in 6-well plates. Cells were cultured in DMEM, 10% FCS, 1% Pen/Strep in a humidified 37°C, 5% CO_2_ incubator until confluent. Media was then supplemented with 50 μg/ml ascorbic acid with complete media changes every 2 days for 20 days. After washing in PBS, cells were denuded by adding Extraction Solution (20 mM NH_4_OH, 0.5% Triton X-100 in PBS) to lyse the cells. Following two washes in PBS, DNA was digested with 10 μg/ml DNase I (Roche) in PBS for 30 minutes at 37°C, before washing twice more. CDMs in tissue culture plates were scraped into 2 × LDS Buffer with reducing agent added before denaturing at 90°C for 20 minutes. CDMs on coverslips were fixed in 10% formalin for 15 minutes at room temperature, washed with PBS and blocked in 30% DS in PBS. Primary antibodies (1:250 Anti-Fibronectin-1, F3648, Sigma-Aldrich; 1:100 Goat Anti-Type I Collagen, 1310-01, Southern Biotech; 1:100 Goat Anti-Type V Collagen, 1350-01, Southern Biotech) were diluted in 30% DS in PBS and incubated on CDMs for 1 hour at room temperature. After 3 washes in PBS, Alexa Fluor conjugated secondary antibodies (Donkey Anti-Goat Alexa Fluor 488, A-11055, Thermo Fisher Scientific; Donkey Anti-Rabbit Alexa Fluor 594, A21207, Thermo Fisher Scientific) were added for 1 hour at room temperature. Finally, 3 washes in PBS and one in water preceded mounting of the coverslips on slides using Vectashield Antifade mounting medium (Vector Laboratories). Imaging was performed using a Nikon Confocal A1R microscope. NIS-Elements or ImageJ software was used for image processing and analysis.

### Statistics

Statistically significant differences between experiments were determined using unpaired Student’s T-test (two-tailed), one-way ANOVA or Welch’s ANOVA test where variances between groups differed significantly. Post hoc analysis using Tukey’s or Dunnett’s multiple comparisons tests was performed (GraphPad Prism v9.1.0). A p-value of less than 0.05 was considered significant. Experiments were performed a minimum of 3 times, unless indicated otherwise, and data is presented as mean ± standard error of the mean (s.e.m.) or ± standard deviation (s.d.).

## Acknowledgements

The authors would like to thank Institute of Genetics and Cancer (IGC) scientific support services, Craig Nicol for slit lamp photography, the IGC Advanced Imaging Resource for help with imaging, the IGC Histology Facility for help with immunohistochemistry, and Edinburgh University Bioresearch and Veterinary Services for animal husbandry. We would also like to thank Stephen Mitchell (University of Edinburgh Biology scanning EM (BioSEM) facility) for assistance with TEM, Toby Hurd for helpful guidance with CRISPR-Cas9 genome editing in mice, and Wendy Bickmore for support and helpful comments on the manuscript.

## Data availability

The datasets supporting the conclusions of this article are included within the article and supplementary files.

## Author contributions

Conceptualization: V.V., C.M.S.; Methodology: C.M.S., A.S,F., V.V.; Software: C.M.S., A.S.F. P.G., V.V.; Validation: C.M.S., A.S.F., V.V.; Formal analysis: C.M.S., A.S.F., C.D., P.G., V.V.; Investigation: C.M.S., A.S.F., C.D., M.Z.M., L.M., P.G.; Resources: C.M.S., A.S.F., V.V., I.J.J.; Data Curation: C.M.S., A.S.F., V.V., Writing - original draft: C.M.S..; Writing - review & editing: A.S.F., I.J.J., V.V.; Visualization: C.M.S., A.S.F.; Supervision: V.V.; Project administration: C.M.S., A.S.F., V.V.; Funding acquisition: V.V.

