## Supplementary Figure 1 for "A Mouse Model of Brittle Cornea Syndrome caused by mutation in *Zfp469*"

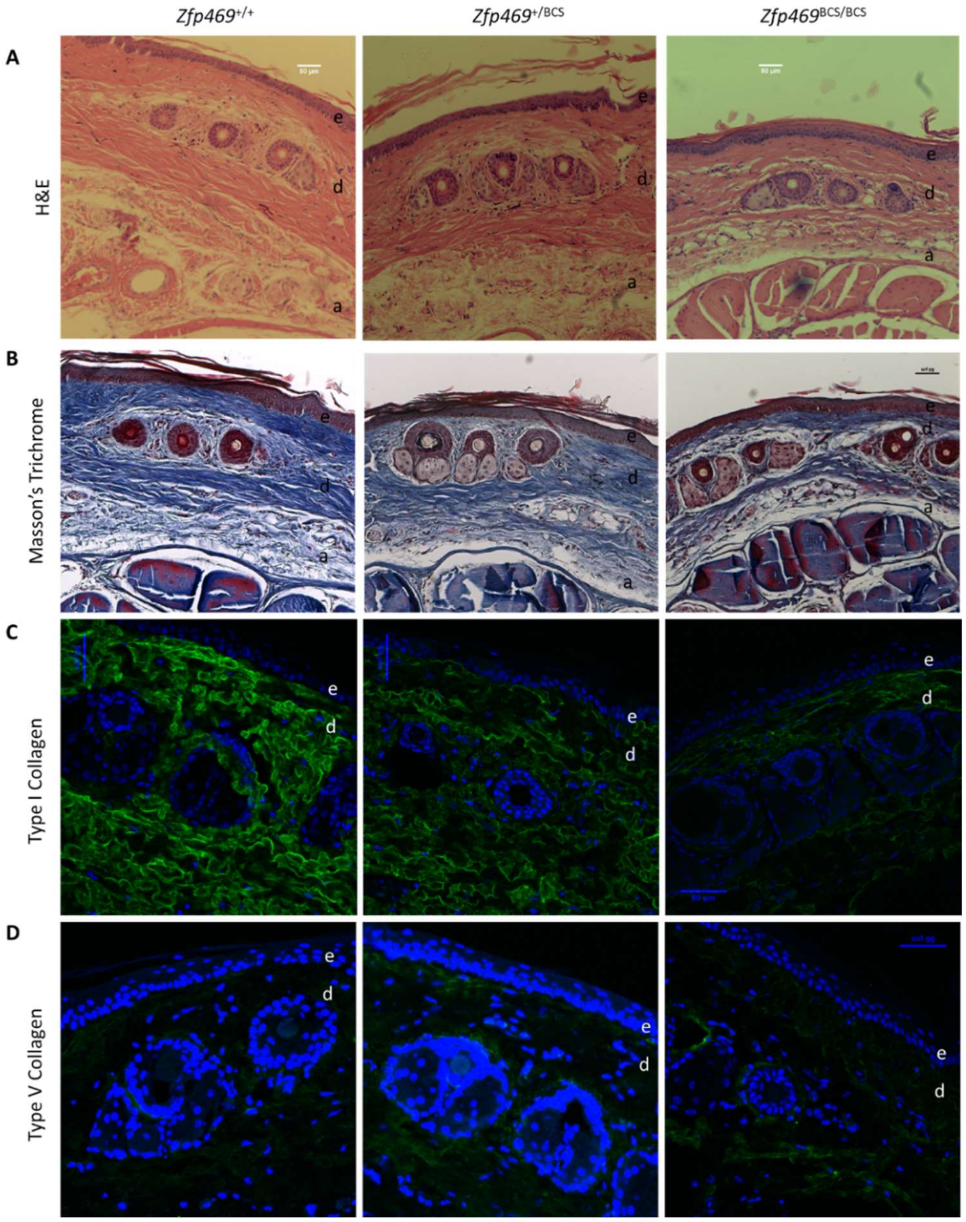


**Fig. S1. Decreased dermal thickness and type I collagen abundance in *Zfp469*$^{BCS/BCS}$ tail skin.** (A) Representative Hematoxylin and eosin (H&E) and (B) Masson's trichrome staining of sections of tail skin from *Zfp469*$^{+/+}$, *Zfp469*$^{+/BCS}$ and *Zfp469*$^{BCS/BCS}$ littermate male mice at 6 months of age shows thinning of the dermis (d) and subcutaneous adipose (a) layer in homozygotes (n=3 for each genotype). The collagen enriched dermal matrix is stained blue by Masson's Trichome stain, the epidermis (e) is stained red. Scale bars represent 50 µm. (C) Representative immunofluorescence images of tail skin sections stained for type I collagen (green) and (D) type V collagen (green) show decreased type I collagen staining in the dermis (d) of *Zfp469*$^{BCS/BCS}$ tail sections, but type V collagen staining is similar to that observed in wildtype tail sections (n=3 for each genotype). Nuclei are stained with DAPI (blue). Scale bars represent 50 µm.
