## Supplementary Table 1 for "A Mouse Model of Brittle Cornea Syndrome caused by mutation in *Zfp469*"

**Table S1. CRISPR-Cas9n genome editing sgRNA and repair template sequences and primers used for in vitro transcription.** Guide RNA sequences shown in bold were used in CRISPR-Cas9 genome editing to generate the *Zfp469*$^{BCS}$ line.

| **Name** | **Sequence 5' - 3'** |
|---|---|
| gRNA1 | TTGAAGGCATCCTCAGCCCCTGG |
| gRNA2 | CTCCCACAAGAGCCCCTCACTGG |
| **gRNA1** | **TTGAAGGCATCCTCAGCCCCTGG** |
| **gRNA3** | **TCCCACAAGAGCCCCTCACTGGG** |
| gRNA4 | AGGGAAGGCTTTGGCTGTCTCGG |
| gRNA5 | TGAGGATGCCTTCAAGAGCCAGG |
| Repair template | CCACACGCTATCAATCCGAGACAGCCAAAGCCTTCCCTCTCCCCACAGAGGGACCA GGCAAACCGATTCCGAACCCGCTGCTGGGCCTGGATAGCACCGGCAAACCGATTCC GAACCCGCTGCTGGGCCTGGATAGCACCTGAGtGCAGAGACGGGTTGAAGGGCTT TCCTCCAGAGCCACCACCTCCACCGCCACC |
| mZNF469g1 T7 primer | TGTAATACGACTCACTATAGGttgaaggcatcctcagcccc |
| mZNF469g3 T7 primer | TGTAATACGACTCACTATAGGtcccacaagagcccctcact |
| Universal reverse | AAAAGCACCGACTCGGTGCC |
