## Supplementary Table 2 for "A Mouse Model of Brittle Cornea Syndrome caused by mutation in *Zfp469*"

**Table S2. Loss of function mutation of Zfp469 is transmitted in the expected ratio in litters born from heterozygote x heterozygote crosses.**

| | +/+ (N) | +/BCS (N) | BCS/BCS (N) | Total (N) |
|---|---|---|---|---|
| **Male** | 8 | 13 | 5 | 26 |
| **Female** | 11 | 23 | 10 | 44 |
| **Obs** | 19 | 36 | 15 | 70 |
| **Exp** | 17.5 | 35 | 17.5 | |

Genotypes of offspring obtained in 10 litters from heterozygote x heterozygote crosses for the *Zfp469*$^{BCS}$ line were subject to Chi2 test, with no significant difference in genotype frequency between observed and expected numbers (N) (two-tailed p value = 0.3787).
